# Nanometer-scale views of visual cortex reveal anatomical features of primary cilia poised to detect synaptic spillover

**DOI:** 10.1101/2023.10.31.564838

**Authors:** Carolyn M. Ott, Russel Torres, Tung-Sheng Kuan, Aaron Kuan, JoAnn Buchanan, Leila Elabbady, Sharmishtaa Seshamani, Agnes L. Bodor, Forrest Collman, Davi D. Bock, Wei Chung Lee, Nuno Maçarico da Costa, Jennifer Lippincott-Schwartz

**Author notes:** Co-first Authors (Carolyn M. Ott, Russel Torres).

## Abstract

A primary cilium is a thin membrane-bound extension off a cell surface that contains receptors for perceiving and transmitting signals that modulate cell state and activity. While many cell types have a primary cilium, little is known about primary cilia in the brain, where they are less accessible than cilia on cultured cells or epithelial tissues and protrude from cell bodies into a deep, dense network of glial and neuronal processes. Here, we investigated cilia frequency, internal structure, shape, and position in large, high-resolution transmission electron microscopy volumes of mouse primary visual cortex. Cilia extended from the cell bodies of nearly all excitatory and inhibitory neurons, astrocytes, and oligodendrocyte precursor cells (OPCs), but were absent from oligodendrocytes and microglia. Structural comparisons revealed that the membrane structure at the base of the cilium and the microtubule organization differed between neurons and glia. OPC cilia were distinct in that they were the shortest and contained pervasive internal vesicles only occasionally observed in neuron and astrocyte cilia. Investigating cilia-proximal features revealed that many cilia were directly adjacent to synapses, suggesting cilia are well poised to encounter locally released signaling molecules. Cilia proximity to synapses was random, not enriched, in the synapse-rich neuropil. The internal anatomy, including microtubule changes and centriole location, defined key structural features including cilium placement and shape. Together, the anatomical insights both within and around neuron and glia cilia provide new insights into cilia formation and function across cell types in the brain.

## Introduction

Primary cilia are membrane enveloped extension of microtubules that grow from the mother centriole in diverse mammalian cells. Because cilia have been difficult to locate in conventional electron microscopy (EM) volumes, we have little knowledge of the structure of cilia in the brain and how the unique extracellular environment of the brain impacts cilia exposure to potential ligands. Although early observers hypothesized a sensory role for neuronal cilia (Allen, 1965; Barnes, 1961), cilia were considered “vestigial and functionless” (Peters et al., 1991) until the surprising observation that induced loss of cilia in adult mice led to increased food consumption (Davenport et al., 2007). Consequently, both the presence and significance of primary cilia in the brain was overlooked by most neurobiologists for generations. (Davenport et al., 2007) Signaling molecules detected and transmitted by cilia are now known to influence brain function, development, behavior, mood, and cognition (Higginbotham et al., 2012, 2013; Koemeter-Cox et al., 2014; Tong et al., 2014; Vien et al., 2023; Guo et al., 2019). Several syndromes caused by defects in primary cilia formation and function include symptoms of impaired physical and cognitive development (Valente et al., 2014), and emerging research suggests links between primary cilia and mental health and age-related decline (Ma et al., 2022; Pruski and Lang, 2019).

Primary cilia share common features across all cell types. The ciliary membrane, continuous with the cellular plasma membrane, sheaths the microtubules of the ciliary axoneme which is anchored to the mother centriole (also called a basal body when a cilium is present). The specialized composition of both the ciliary membrane and cytoplasm is maintained by the transition zone, a boundary formed by repeating “Y”-shaped protein complexes that ring the microtubules at the base of the cilium (Park and Leroux, 2022). Cilia microtubule doublets extend directly from mother centriole microtubules. Despite the stereotyped structure of cilia, both classic and emerging data support cell specific differences in cilia structures in different systems and tissues (Soares et al., 2019; Doroquez et al., 2014). For example, although typically illustrated as extending the entire length of the cilium, microtubule doublets can change and even terminate within a cilium (Kiesel et al., 2020; Klena and Pigino, 2022). In the visual cortex the different types of neurons and glia have been grouped into cell classes: excitatory neurons or inhibitory neurons and astrocytes, microglia, oligodendrocytes, or oligodendrocyte precursor cells (OPCs). Given the strong links between primary cilia and brain health, we asked whether cilia exhibit structural differences relevant for their functioning in these different cell classes.

Early efforts at electron microscopy captured serial sections of a few astrocyte and neuron cilia (Allen, 1965; Dahl, 1963; Herman et al., 1971; Spacek, 1985). However, cell-specific structural knowledge in the context of neural tissue has been elusive because primary cilia have been difficult to locate using conventional electron microscopy (EM) approaches. There is typically only a single cilium per cell and, upon exiting the cell, the cilium is buried in the network of glial and neuronal processes, called the neuropil. This meshwork is the context for non-synaptic signal transmission to neuron and glia cilia. A growing number of immense EM volumes are being generated, motivated by efforts to locate synapses and trace network connectivity and the high-resolution datasets include hundreds or thousands of cells in brain tissue. In addition, modern computational innovations used to generate 3D volumes of each cell body and process have provided unprecedented access to multiple quantifiable cellular dimensions. Thus, an opportunity now exists for employing these datasets for gaining unique views of cilia and other cellular structures in the diverse cell types of the brain.

Here, we mine three 3D transmission electron microscopy (TEM) volumes of the mouse primary visual cortex (Bock et al., 2011; Schneider-Mizell et al., 2021; Dorkenwald et al., 2022; Turner et al., 2022; Buchanan et al., 2022; Elabbady et al., 2022) to determine primary cilia ultrastructure in hundreds of cells across different brain cell classes. We discovered key anatomical differences between cilia on neurons and glia, including differences in microtubule organization and basal body docking. In addition, we re-image selected cilia at higher resolution to resolve subtle differences between excitatory neuron and astrocyte transition zones. We characterized, for the first time, the dense extracellular micro-environment surrounding primary cilia and found that cilia pass directly adjacent to synapses. Synapses are not enriched near cilia, rather, cilia stochastically encounter synapses that densely populate the surrounding neuropil. We further analyzed physical parameters of cilia, such as shape and orientation, and demonstrated cell-class differences in cilia structure and formation. Together, the new data reveal variations in primary cilia anatomy in the brain and indicate that cell-type specialization of cilia structures may influence brain development, behavior, mood, and cognition.

## Results

### Locating cilia in neurons and glia in the visual cortex

The generation of EM volumes capturing hundreds of cells in mouse visual cortex created an opportunity to characterize primary cilia in their native context. To do this we located and annotated cilia and centrosomes in three volumes, which we refer to in the text based on the age of the mouse: P36, P54 and P>270. In addition to animal age, the datasets also include different layers of visual cortex as described in Sup. Table 1. Figure 1A shows cellular reconstructions of representative cells from five morphologically defined cell classes in the primary visual cortex (V1) EM volumes (i.e., excitatory neurons, inhibitory neurons, astrocytes, OPCs, oligodendrocytes and microglia) based on volumetric EM image segmentation. The most prevalent cells were excitatory pyramidal neurons. Several types of inhibitory neurons were observed: including chandelier cells, basket cells, neurogliaform cells, bipolar cells, and Martinotti cells. Glial cells that were observed included astrocytes (both fibrous and protoplasmic), microglia and cells in the oligodendrocyte lineage, including precursor cells (OPCs), premyelinating and mature oligodendrocytes.

**Figure 1:**
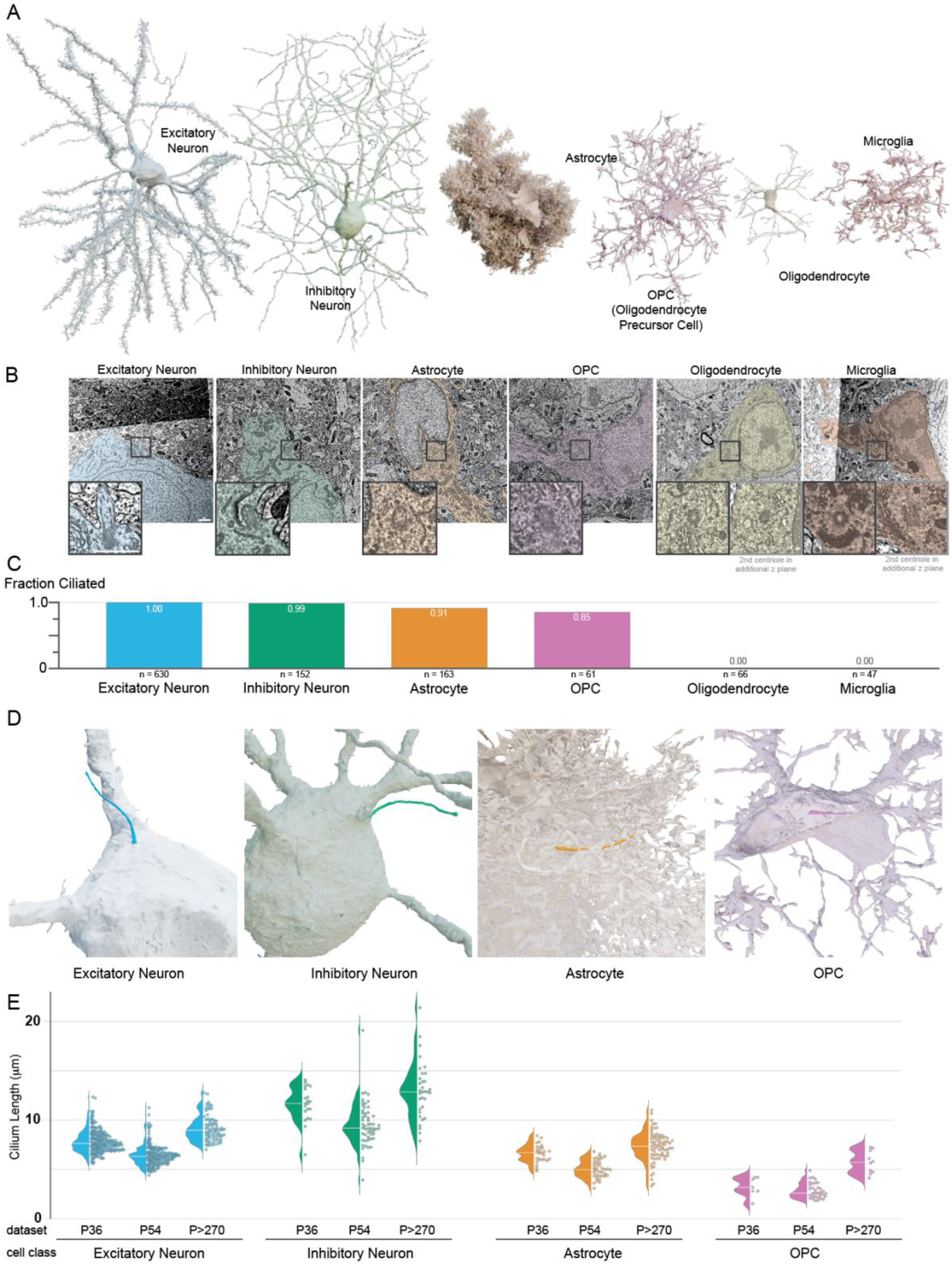
Primary cilia are pervasive on neurons, astrocytes and OPCs in primary visual cortex, and absent from oligodendrocytes and microglia. **A.** 3D renderings of the six cell classes in primary visual cortex annotated in this study. **B.** Electron micrographs of the cells rendered in A. The inset images show either the base of the cilium or both centrioles for cells that are nonciliated. Scale bar in both image and inset is 1 μm. **B.** Plot of the fraction of cells with cilia by cell class. **D.** 3D renderings of the cilia on the cells shown in A. Portions of the astrocyte and OPC were cut away to reveal the cilium. **E.** Cilium length of every cilium annotated is graphed both as a distribution (left) and individual value (right).

To assess which of these cells had primary cilia, we annotated the position of both the mother and daughter centriole in each cell. Figure 1B shows the EM images that correspond to the base of the cilium for the two neurons, the astrocyte and the OPC cells displayed in Figure 1A. Both centrioles are included for the oligodendrocyte or microglia, neither of which were ciliated. The extent of ciliation for each cell class is shown in Figure 1C and detailed in Table 1. We found cilia on all excitatory neurons and on 150 out of 152 inhibitory neurons. In addition, 91% of astrocytes (148 of 163) and 85% of OPCs (52 of 61) were ciliated (Buchanan et al., 2022). (Of the 9 nonciliated OPCs, 8 had centrosome associated ciliary vesicles, which will be discussed later). No cilia were found on any of the 66 oligodendrocytes or 47 microglia assessed, which is consistent with other brain regions (Sipos et al., 2018; Falcón-Urrutia et al., 2015; Bear and Caspary, 2023). Figure 1D shows examples of cilia on an excitatory neuron, inhibitory neuron, astrocyte and OPC, with the meshwork of surrounding glial cell body and/or processes that obscure cilia removed.

**Table 1.**

| **Cell Class** | **P36 \*\***<br>**% ciliated**<br>**(ciliated/assessed)** | **P54#**<br>**% ciliated**<br>**(ciliated/assessed)** | **P>270**<br>**% ciliated**<br>**(ciliated/assessed)** | **Total cells**<br>**annotated** | **Total %**<br>**ciliated** |
| --- | --- | --- | --- | --- | --- |
| Excitatory | 100%<br>(292/292) | 100%<br>(256/256) | 100%<br>(82/82) | 630 | 100% |
| Inhibitory | 100%<br>(32/32) | 99%<br>(85/86) | 97%<br>(33/34) | 152 | 99% |
| Astrocyte | 95%<br>(35/37) | 78%<br>(42/54) | 100%<br>(71/72) | 163 | 91% |
| OPC | 67%<br>(8/12) | 87%<br>(34/39) | 100%<br>(10/10) | 61 | 85% |
| Oligodendrocyte | 0%<br>(0/2) | 0%<br>(0/57) | 0%<br>(0/7) | 66 | 0% |
| Microglia | 0%<br>(0/10) | 0%<br>(0/22) | 0%<br>(0/15) | 47 | 0% |

\* All cells in the volume were assessed

A key feature of primary cilia is length because cilia length determines how far a cilium extends into the surrounding environment, the surface area available for signal detection, and the volume of the internal signaling compartment. To compare cilia length within and between cell classes, we first generated a skeleton line that tracks the central path from the base to the tip, either by manually tracing the cilium or by calculating the central path of the segmented structure. We then measured the length of each cilium. The measured cilia pathlengths are plotted by cell class for each dataset in Figure 1E. While all three datasets encompass the V1 visual cortex, they are not replicates. Overall, the inhibitory neuron cilia were the longest and OPC cilia were the shortest. Within each dataset, the average astrocyte cilium length was less than the average excitatory neuron cilium length. We found that while cilia lengths clustered for each cell type, there were differences between the datasets. Although cilia in P>270 generally appear longer than in P36 or P54 (See Figure 1E and Figure S1A) differences in sample preparation could produce minor discrepancies that affect scaling. As highlighted in Sup Table 1, each dataset included a unique sampling of visual cortex layers, which also contributed to differences between the datasets. For example, the average inhibitory neuron cilium length in P36, which contains mostly layers 2/3 and a portion of layer 4, was 11.6 μm +- 0.7 μm (95% ci). The average inhibitory neuron cilia length in layer 2/3 of P54 was comparable to P36 (11.6 μm +- 1.8 μm (95% ci), however, the average length for the entire volume was less (9.6 μm +/-0.6 μm; 95% ci).

In addition to comparing cilia length between cell classes, we were also able to compare cilia lengths between cell types within some classes. Most astrocytes in the datasets were protoplasmic astrocytes, however, the P>270 volume includes 20 fibrous astrocytes – many of which are in Layer 1. We found that the cilia of fibrous astrocytes were shorter on average than the protoplasmic astrocytes in the P>270 volume (Figure S1A). The inhibitory cell class included many different cell types, some of which have been specifically identified in the P36 and P54 volume. The differences between cilia lengths for the identified basket cells, bipolar cells, chandelier cells, Martinotti cells and neurogliaform cells shown in Figure S1B suggest that cilia in individual cell types cluster more strongly than the cell class as a whole. Different types of inhibitory neurons populate the layers of the cortex (Kim et al., 2017), so both the observed differences in average cilia length mentioned above and the distributions of cilia lengths (plotted by depth in Figure S1C) are indicative of differences within the inhibitory cell class. For excitatory neurons we were able to uncover potential cell type differences in cilia length by comparing cilia across the layers of the P54 volume. As shown in Figure S1C, we observe a larger variance in cilia length in Layer 5 than in the other cortical layers. While some layer 5 cilia are close to the average, many are among the longest excitatory cilia. Together, the cilia length measurements revealed differences in cilia length between cell classes and within cell classes, which may impact cilia exposure to external signals.

### Ciliary pockets are found predominantly in astrocyte and OPC cells

To understand the physical properties of cilia in the cortex, we began by leveraging the resolution of the EM volumes to examine and compare structural features of cilia. We observed differences in the origin of the cilium between cell classes. Specifically, in some cells the cilium extended directly from a centriole docked at the plasma membrane (surface cilium; Figure 2A). In others, the cilium extended from a centriole recessed in the cytoplasm that had a pocket of membrane that dipped from the surface to anchor at the distal appendages (pocket cilium; Figure 2B). The form of an individual cilium is thought to be determined by its biogenesis pathway (Zhao et al., 2022; Sorokin, 1968). A surface cilium grows from a mother centriole that has docked at the plasma membrane and the entire length remains externally exposed. In contrast, a pocket cilium originates as a membrane-enclosed internal structure that retains the encapsulating membrane to form the ciliary pocket.

**Figure 2:**
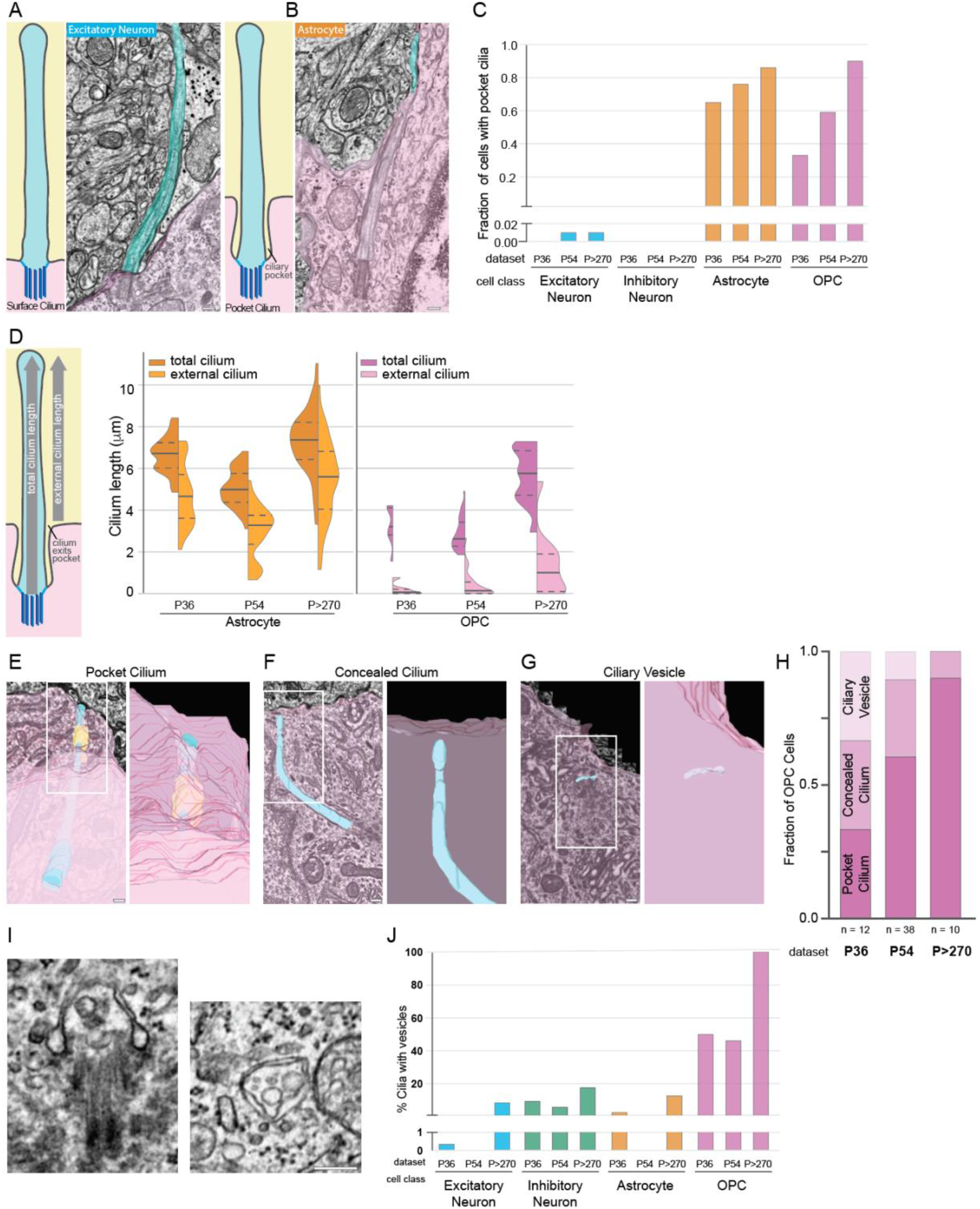
Neuron cilia dock directly at the plasma membrane, while glia cilia are associated with a ciliary pocket. **A - B.** Illustrations of cilia either docked directly at the plasma membrane (Surface Cilium) or recessed in a ciliary pocket (Pocket Cilium) are paired with 3D segmented representations of (A) an excitatory neuron and (B) an astrocyte cilium (cyan) and the cell interior (pink) overlaid onto a single EM plane. **C.** For each cell class from each EM volume, the fraction of cilia emerging from a ciliary pocket is graphed. **D.** The ciliary pocket shields the base of the cilium so only a portion of the cilium is external. The distribution of cilium lengths for astrocytes and OPCs in each volume is plotted adjacent to the distribution of the external length of the cilium. **E - G.** 3D segmented OPC cilia (E, F) or ciliary vesicle (G) (cyan) and cytoplasm (pink) overlaid onto EM images. The ROIs are enlarged to the right of each image to illustrate that the cilium barely emerges (E) or is completely concealed (F). All OPCs without cilia had ciliary vesicles associated with the mother centriole (G). (Several OPC have membrane structures in the ciliary pocket illustrated here in yellow.) Scale bar is 200 nm; inset ROI has a length of 1085 nm. **H.** The fraction of cilia of each classification is plotted for each volume. *n* is the number of OPC cells examined in each volume. **I.** Example images of vesicles inside OPC primary cilia. Scale bar is 200 nm. **J.** The percentage of annotated cilia with internal cilia vesicles was graphed for each cell class in each dataset.

To assess the differences between cell classes, we quantified the fraction of cells with a ciliary pocket. As shown in Figure 2C, with rare exceptions all neurons had surface cilia. In contrast, glial populations predominantly had pocket cilia. Because the ciliary pocket limits the external exposure of cilia, we quantified the pathlength of the external portion of the cilium in cells with pocket cilia. The total pathlengths and external pathlengths for astrocyte and OPC cilia from each dataset are plotted Figure 2D. Generally, the external length of astrocyte cilia was ∼2 μm shorter than the total length. OPC cilia, which are shorter than astrocyte cilia, had an even larger difference between the exposed length and total length. The majority of OPC cilia project less than 3 μm from the cell – in P36 and P54 that number was even less. The fraction of the cilium shielded by the ciliary pocket was dependent on both the depth of the pocket and the total length of the cilium. OPC cilia were generally the shortest cilia observed in these datasets and in many OPC cells the cilium only extended 1-2 μm into the surrounding neuropil. These data indicate that astrocyte and OPC cilia have less exposed signal detecting surface area than neuronal cilia.

Most of the astrocytes that don’t have pocket cilia have surface cilia. The same is not true for OPCs. Instead, we noticed that OPCs without pocket cilia had cilia structures that indicated they were in the process of ciliogenesis or cilia disassembly. To examine how these intermediate structures compare across datasets, we classified them as either concealed cilia (cilia that are entirely contained in the cytoplasm and do not exit) or ciliary vesicles (mother centrioles with membrane structures associated with the distal appendages). Figures 2F, 2G and 2H present examples of each with an accompanying 3D segmentation. All concealed cilia and ciliary vesicles were found in OPC cells with one exception (an astrocyte daughter centriole in P>270 had a ciliary vesicle). The fraction of OPC cells with pocket cilia, concealed cilia and ciliary vesicles in each dataset are graphed in Figure 2H. We observed a trend that correlates with the ages of the mice. OPCs in the P36 dataset had pocket cilia, concealed cilia and cells with ciliary vesicles. The P54 dataset had a higher fraction of pocket cilium, a similar fraction of concealed cilia and fewer cells with a ciliary vesicle. In the oldest animal all OPCs except one had pocket cilia.

Within the ciliary pocket of several OPC cells, we found inclusions (shown in yellow in Fig 2E). It is possible that these large, membrane-enclosed vesicles were derived from evagination of the ciliary membrane or the ciliary sheath (Wang et al., 2021a; Wood et al., 2013; Wood and Rosenbaum, 2015). Alternatively, these could be portions of neuronal or astrocytic processes that have pinched off. The latter possibility is suggested by the presence of such processes in the ciliary pocket of some cells. We also observe inclusions in the pocket of submerged cilia. If the source of the inclusion were neuronal or astrocytic processes, it would suggest that emergence of the cilium at the cell surface could be reversible, a process which has been proposed in cultured fibroblasts (Rivera-Molina et al., 2021).

An unanticipated observation revealed by EM was the presence of vesicles inside cilia (Figure 2 I, 2J and Figure S2). Although infrequent, there are published TEM images of vesicles inside primary cilia (Arellano et al., 2012; Reese, 1965; Banks et al., 2017; Wakefield and Waite, 1980; Shah et al., 2008). In astrocyte and neuron cilia the observed vesicles were rare. When observed in astrocyte and neuronal cilia, we typically found a single vesicle, however, occasionally two or more clustered together (Figure S2A). In contrast to a previous suggestion that vesicles could move along the center of the microtubules, all vesicles were adjacent to the axoneme microtubules (Ruba et al., 2023). Vesicles inside OPC cilia were prevalent and diverse (Figure S2B). We found single-membrane, round vesicles with transparent contents. We also observed single-membrane compressed vesicles, vesicles with dark centers, and double-membrane vesicles (Figure S2B). As our observations are from a single time point it is difficult to discriminate whether the vesicles originate in the cell body and are transported into the cilium for exocytosis, or if they are generated by endocytosis of the ciliary membrane. Together, the analysis of cilia nanostructure reveals several distinctions between cell classes.

### The transition zone in neuronal cilia persists farther than in astrocyte cilia

We also examined the boundary structure at the base of the cilium: the transition zone. While both the cytoplasm and membrane of cilia are continuous with the rest of the cell, the transition zone creates a boundary which both facilitates selective exclusion and promotes active retention of cilia-localized proteins. When sectioned along the cilium’s length, transition zones appear as a density between the ciliary membrane and the microtubules (Figure 3A, B). In cross-sections, the structural components of the transition zone make a “Y” shaped structure referred to as a Y-link (Gilula and Satir, 1972) (Figure 3C and Figure S3). Internal structures were especially clear in the P>270 dataset and our initial impression during annotation was that astrocyte transition zones might be reduced compared to neurons, but this was challenging to assess both because the structure was not fully resolved and because many cilia are sectioned at obtuse angles. We used two strategies to try to discriminate differences between neuron and astrocyte transition zones. To investigate structural details at higher resolution we selected seven cilia in the P>270 dataset found to be primarily aligned either parallel or perpendicular to the imaging plane and reimaged these cilia on the original TEM grids at higher resolution, ranging from 0.36 to 0.56 nm per pixel (compared to 4 nm per pixel for the original dataset), and registered the high-resolution reimaging to the original image volume. Figure 3A illustrates the improvement in resolution achieved by re-imaging the same cilium at higher resolution (4 nm (left) and 0.56 nm (right) per pixel XY resolution). In Figure 3B the transition zones of an excitatory neuron and astrocyte cilium are visible in alternating sections due to the periodic nature of the structure. The transition zone density between the cilium membrane and the microtubules in several of the z sections (Figure 3B) extends farther in the neuronal cilium than in the astrocyte cilium. Y-links were visible in both astrocytes and neurons (Figure 3C); however, they were visible in fewer sections of the astrocyte cilium (Figure S3A and S3B). Another distinction between transition zones was only visible in fibrous astrocytes: we observed electron dense materials filling the center of the cilium (Figure S3C). These cells have intense intermediate filament staining throughout the cell body and it seems possible that some filaments extend inside the cilium. The internal staining cleared in the medial and distal portions of the cilium.

**Figure 3:**
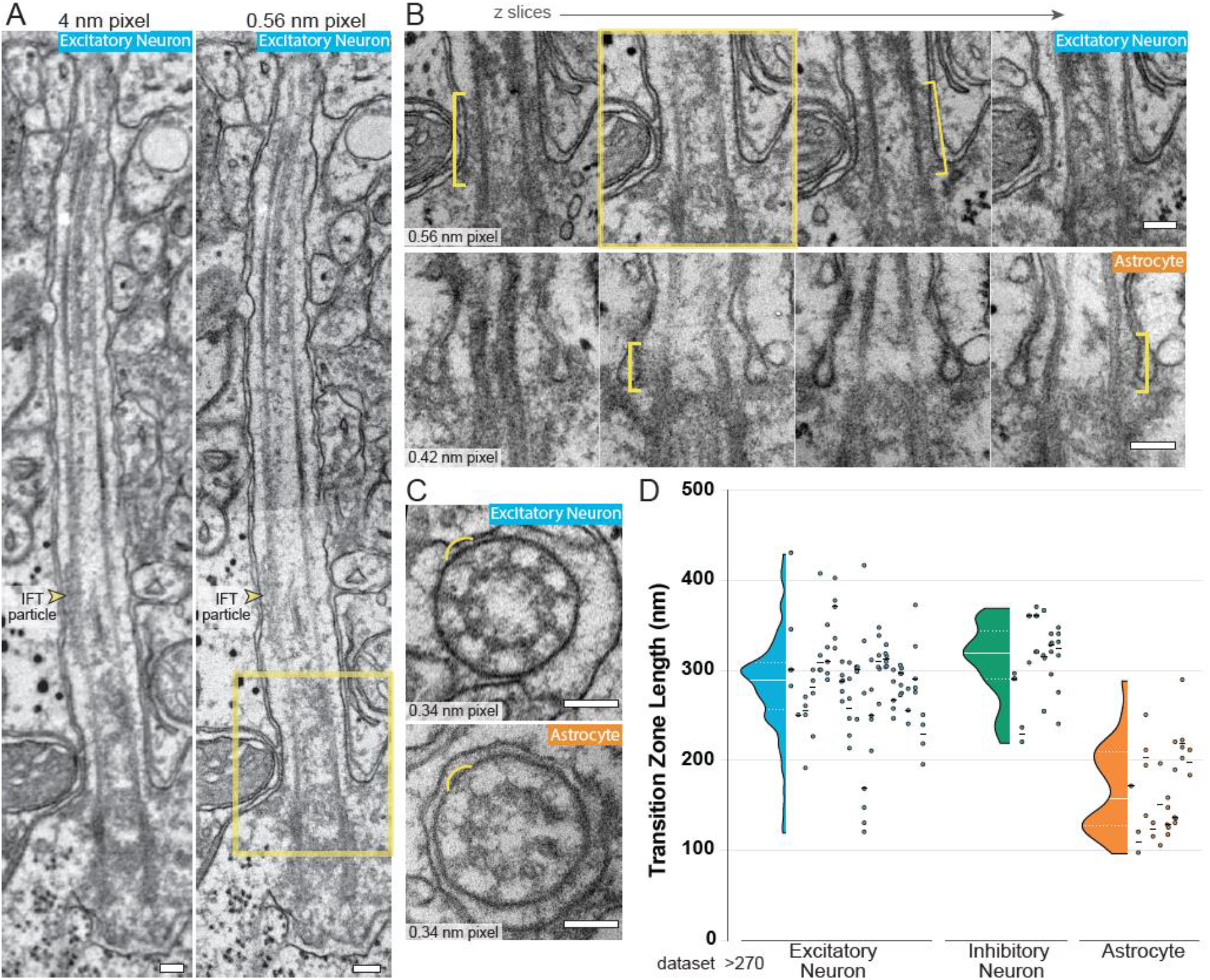
Astrocyte transition zones are shorter than neuron transition zones. **A.** To improve the resolution of the transition zone, specific cilia from the P>270 dataset were located on the original EM grids and reimaged at higher resolutions. In A, the image on the left is from the original dataset; the panel on the right was reimaged at higher resolution (two images were aligned to show a similar area). An IFT particle is visible between the ciliary membrane and a microtubule. **B.** Serial images of the base of an excitatory neuron cilium and an astrocyte cilium are shown. The yellow box indicates the image that matches the ROI in A. The transition zone is visible in alternating slices and is highlighted by a yellow bracket. **C.** The expected Y-link structures (yellow arcs) of the transition zone are resolved in high-resolution images of cilia cross-sections in both excitatory neuron and astrocyte cilia. **D.** The length of the transition zone of each cilium sectioned lengthwise in the P>270 dataset was measured in sequential images. The distribution of all length measurements is plotted on the left for each cell class. On the right, the measurements for each individual cilium are plotted in the same column and the mean length for that cilium is represented by a black line. Differences in TZ measurements within a single cilium are likely due to the position of the cilium relative to the sectioning plane. Scale bars in A, B, and C are 100 nm.

The second strategy we used to compare transition zones in astrocyte and neuron cilia was to identify a subset of cilia with the base sectioned approximately parallel to the imaging plane in the P>270 dataset. We then measured the transition zone density in every z plane of each cilium. Figure 3D presents both the individual values with an average for each cilium and composite measurements as a violin plot. Variations in transition zone measurements within a single cilium were expected because the cutting angles were not perfectly parallel. The range of transition zone measurements for both excitatory and inhibitory neurons was noticeably larger than in astrocytes (Figure 3D). The process of transition zone formation has not yet been examined in neurons and astrocytes, but the structural differences observed might result from differences in assembly as demonstrated in other tissues (Wiegering et al., 2018; Akella et al., 2019; Jana et al., 2018).

### Microtubules and microtubule inner binding proteins change from the ciliary base to tip

To further examine differences between cilia in neurons and glia we examined the microtubules within each cilium. It is known that the number and organization of microtubules can change from the base to the tip (Sun et al., 2019; Kiesel et al., 2020; Gluenz et al., 2010). Axoneme doublets include an A-tubule and a B-tubule that closes onto the outside wall of the A-tubule. We noticed several changes in microtubules as they progress from the cilium base to the tip (diagrammed in Figure 4A). The cilia in Figure 4B and C (and in their entirety in Figures S4 and S5 and Sup Video 2) illustrate these changes because these cilia only bent slightly and were oriented perpendicular to the imaging plane, thus providing an optimal view of the microtubules along the entire length. We found that the A-tubules had an electron-dense center near the cilium base. We suspect the densities in the A-tubule are microtubule inner binding proteins. Recent investigations using cryoEM have identified and characterized proteins that bind to the inside of the *Chlamydomonas* flagellar doublet microtubules (Ma et al., 2019; Ichikawa et al., 2019). Beyond the proximal region of the cilium, the center of the A-tubule appeared translucent suggesting that microtubule inner binding proteins are restricted to the base of the cilium. The change in A-tubule staining does not happen to all doublets at the same distance from the centriole. In Figure 4B and C, the doublets tracked with yellow arrow transition approximately 2.3 μm (excitatory neuron) and 1.2 μm (astrocyte) from the base. Although there appears to be a region where the transition occurs, the densities in the A-tubules persist longer in neuronal cilia. The first cleared doublet in the excitatory neuron is in the 25th slice (approximately 1.2 μm from the base; Figure S4) and most doublets appeared cleared after the 43rd slice (∼1.9 μm from the base). In contrast, cleared doublets were visible in the 5^th^ slice of the astrocyte cilium (∼225 nm from the base; Figure S5). In neurons the dark to light tubule transition typically happened once. In contrast, several astrocyte doublets alternated between filled and empty (see doublets #4 and #7 in Figure S5). The changes in microtubule staining can also be seen in cilia cut along the cilium length (Figure S6A). Dark microtubule staining is visible at the base of the cilium where the two sides of the doublet are difficult to resolve in the side view. Beyond the proximal cilium, the staining intensity decreased.

**Figure 4:**
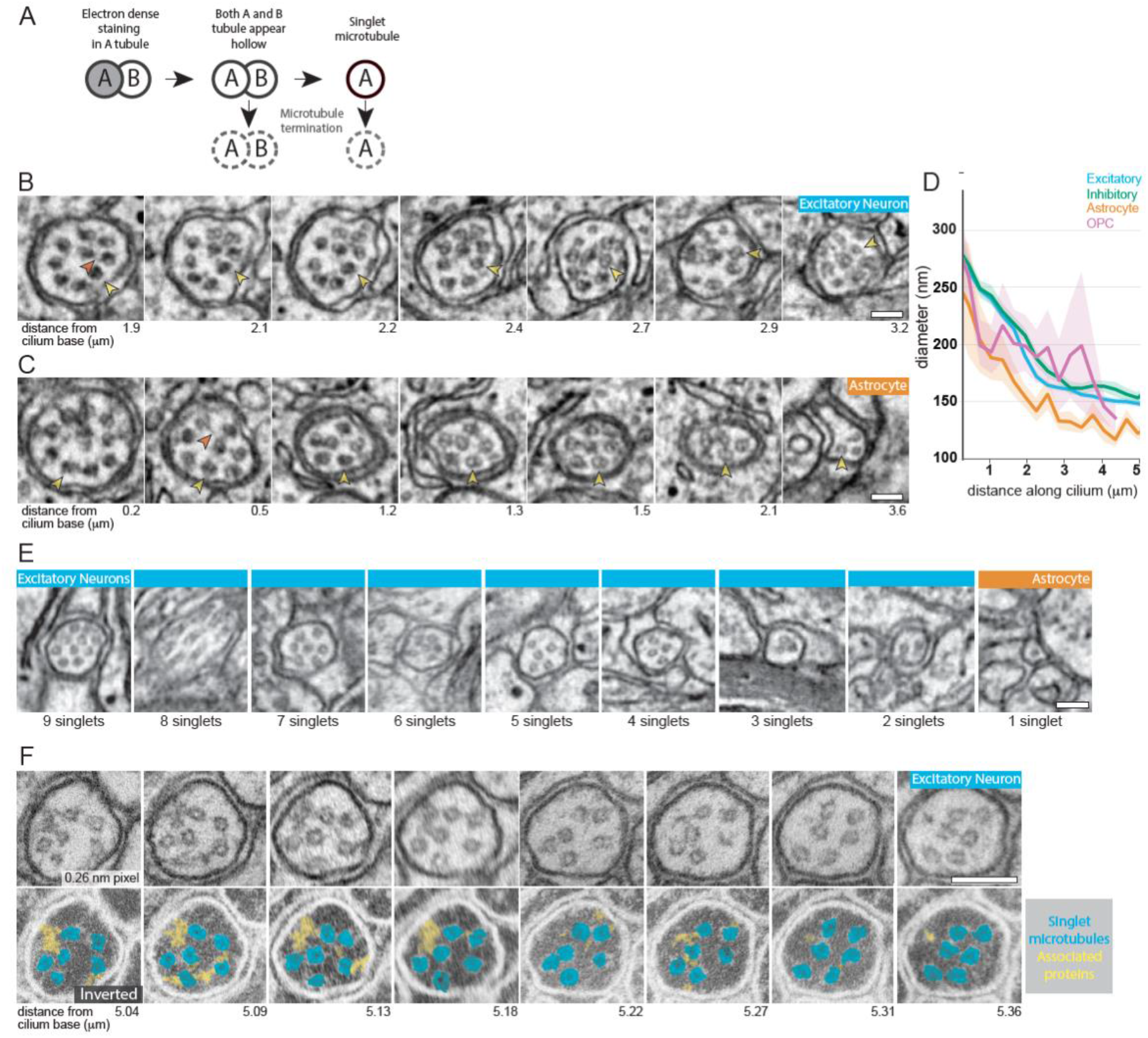
Microtubule structures reconfigure differently within astrocyte and neuron cilia. **A.** The illustration represents the microtubule structural transitions observed within cilia. Near the basal body microtubules emerge as doublets configured with A- and B-tubules. Density within the A-tubule ends leaving translucent lumens in both tubules. Some of the microtubules transition to singlets. Both doublet and singlet microtubules have been observed to terminate. **B - C.** Cross-sections starting near the base of an excitatory neuron cilium (B) and an astrocyte cilium (C). The distance of individual planes from the base of the cilium is indicated below each image (the entire image series are presented in supplemental figures 3 and 4). The yellow arrowheads track the position of the same microtubule through the volume. The orange arrowhead points to densities between the deviant microtubule and adjacent doublets. **D.** The diameter along the central path length of a proximal portion of the cilium was measured for the cilia in the P54 dataset. The solid lines represent the average diameter of a binned pathlength, with the shading representing the 95% confidence interval of that bin. **E.** Cross-sections near the tips of different cilia display the diversity of microtubule configurations. **F.** Serial cross-sections from the distal region of an excitatory neuron cilium were located on the original grids and imaged at higher resolution. The images in the lower panel have been inverted and colored to highlight the electron densities (yellow) that bridge between the ciliary membrane and microtubules (cyan) or between individual microtubules. All scale bars are 100 nm.

Also evident in Figures 4B and C was the movement of a doublet from the periphery into the center of the axoneme in the proximal cilium, shifting the microtubule organization from 9+0 (9 peripheral microtubule doublets) to 8+1 (8 peripheral doublets and a single central doublet) (Allen, 1965). We observed deviant microtubules in all cilia (also shown in Figure S6A or S6C). Occasionally two doublets entered the center (7+2 configuration).

We were also able to document partial and complete microtubule terminations (also illustrated in Figure 4A). Partial termination refers to the termination of B-tubules, which leaves singlet microtubules (yellow arrows in right panels of Figure 4B and Figure S4 and S5) whereas complete termination refers to termination of both A- and B-tubules. In astrocyte cilia, we observed termination of both doublet and singlet microtubules (see doublets #2, #3, and #6 in Figure S5). In neuron cilia the terminating filaments were largely singlet microtubules. In serial cross-sections it was not possible to resolve any structures at the microtubule terminus. We did capture a side view of microtubule termination in an astrocyte cilium while re-imaging the transition zone at high resolution (Figure S6C). Electron densities on either side of the terminating microtubule –one side contacting the ciliary membrane – suggest the presence of a capping or termination structure.

To investigate how the internal changes in cilia microtubules affect the overall structure of the cilium, we measured the diameter along the length of the 3D segmented cilia. All cilia narrow within the first 4-5 μm in the region where microtubule transitions and termination happen (Figure 4D). We found differences between cell classes: the diameters of both inhibitory and excitatory neuron cilia decreased steadily along the first 3 μm. Astrocyte cilia diameters appeared to drop off more quickly. An example of the difference in the rate of diameter change is visible in Sup. Video 2. This is consistent with our observation that microtubules in astrocyte cilia departed from the 9+0 configuration closer to the base than neurons. OPC cilia diameters generally dropped quickly like the astrocyte cilia, however, there is much more variance in diameter between OPC cilia. The prevalence of vesicles inside cilia may contribute to this difference. In addition, the ciliary membrane of OPC cilia does not appear to constrict uniformly as microtubules terminate (example cross sections of OPC cilia are shown in Figure S6E).

To investigate whether termination of microtubules would be stereotyped across cilia of a single cell class or cell type we examined every annotated cilium in the >P270 dataset and quantified the number of singlets present at the most distal resolvable section (although microtubule structure is generally clear in the P>270 dataset insufficient resolution and cut angles prevented quantification in many cilia). All axonemes change along the length and we found examples of every possible number of singlets in the distal portion of cilia (see gallery of examples in Figure 4E). In many cells the cut angle made it difficult to discern the number with confidence. We plotted the number of singlets near the terminus by cell type (Figure S6D) and observed a possible trend that astrocyte and OPC cilia are more likely to have only a few singlets near the tip and neuronal cilia are most likely to have 4-6 singlets.

Little is known about the function or prevalence of microtubule binding proteins in brain cilia. As we examined the microtubules, we found densities that appeared to tether microtubules to the ciliary membrane or to each other. Near the base of the cilium, we observed densities between the axoneme microtubules and the ciliary membrane (Figure S6B). We could also see densities that linked microtubules to the membrane in cross-sections (Figure 4B and Figure S6E). It is possible that these densities are similar to those observed in the kidney cilium and in cultured epithelial cells (Gluenz et al., 2010; Sun et al., 2019). We also observed previously unreported densities that connect the deviant microtubule to adjacent doublets in cross-sections and side views (Figures 4B and Figure S6B and S6C). These bridging structures suggest that the reorganization is accomplished or supported through inter-doublet protein bridges.

To carefully examine microtubules and associated proteins in the distal region of the cilium we reimaged a cross-sectioned, excitatory neuron cilium at higher resolution (Figure 4F). We found evidence for bridging complexes associated with singlet microtubules. In addition, densities created a bridge between microtubule singlets or between a singlet and the ciliary membrane. We observed a density that fills an area between the ciliary membrane and three microtubules in four serial sections. The length suggests it could be an unconventional IFT train. (Conventional IFT trains are thought to run along a single A or B-tubule (Stepanek and Pigino, 2016). The identity and function of the various microtubule associated densities is unclear, however their location and prevalence suggest that they may contribute to axoneme stability.

### Cilia project into a tangle of neural and glial filaments

With an established understanding of the physical properties of cilia we expanded our investigation to the environment outside the cilium, which is invisible in conventional immunofluorescent investigations of cilia. In the TEM volumes we tracked cilia and quantified the types of structures adjacent to neuronal and glial cilia. This was possible in the densely segmented datasets (P36 and P54), where we could visualize and navigate through the 3D data to classify each structure based on origin, shape and identifying features such as pre and postsynaptic densities or glycogen granules. We found that as they emerge many cilia remain proximal to the cell body while others project away directly. Outside the cell, cilia encounter the neuropil, the tangle of local axons, dendrites, and glial processes. Figure 5A and B provide representative views with the complex extraciliary environment with each process shaded by type (the relevant volumes are visible in Sup Video 3 and 4). Colored regions distant from the cilium in this focal plane pass next to the cilium in a different plane. To communicate the complexity of the cilium niche, we generated 3D renderings of each cilium and adjacent axons, dendrites, or glial processes (Figure 5C and 5D). In Figure 5D the portion of the cilium shielded by the ciliary pocket has no adjacent processes.

**Figure 5:**
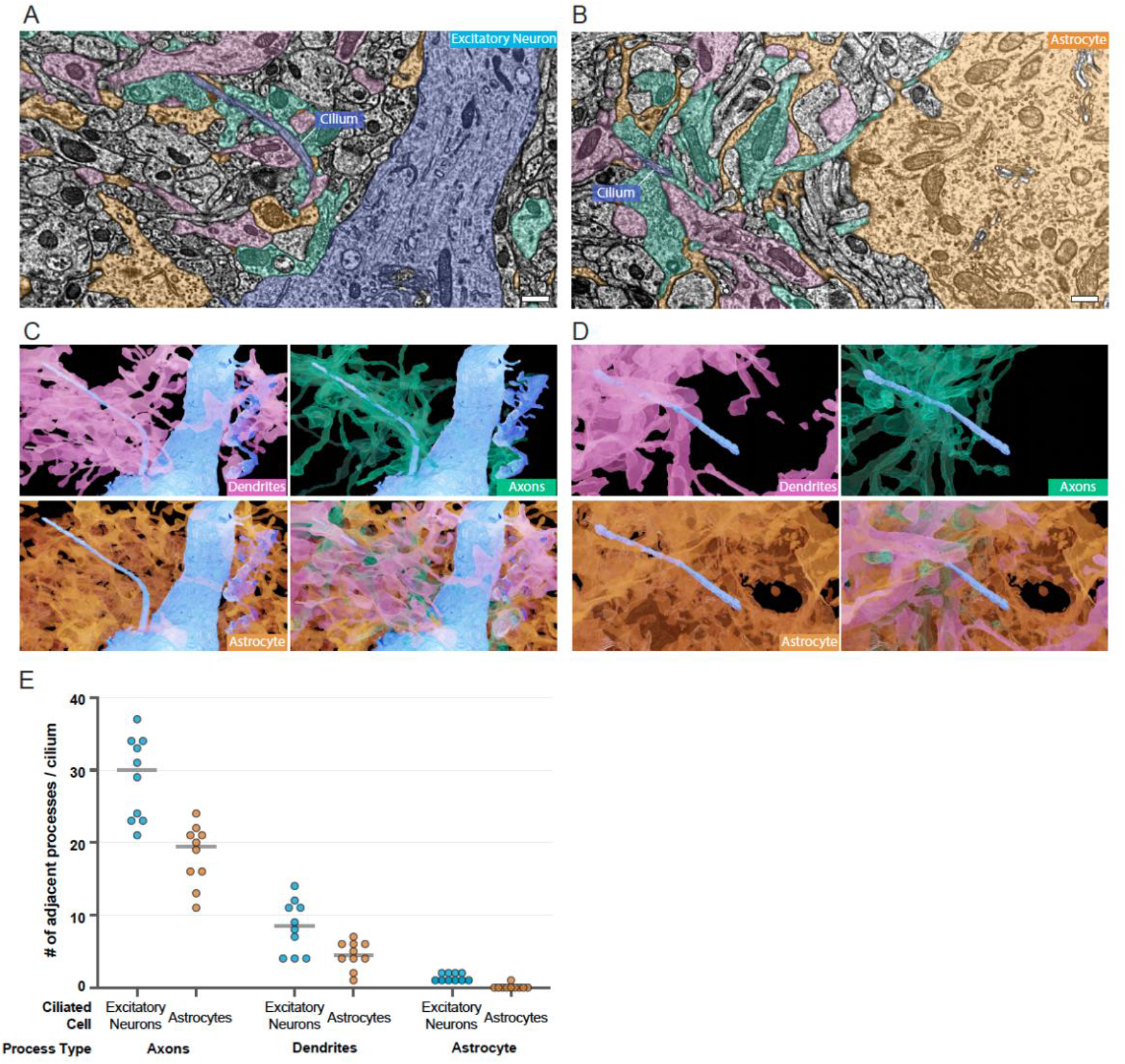
The ciliary pocket reduces the number of processes adjacent to astrocyte cilia in the dense neuropil. **A - B**. Cilia on an excitatory neuron (A) or Astrocyte (B) in the P36 dataset are shown in blue. Every process in the neuropil that passes adjacent to any portion of the cilium was colored by processes type: dendrites are pink, axons are teal and astrocyte processes are orange. The apical dendrite of the excitatory neuron is blue (A), and the cytoplasm of the astrocyte is also orange (B). **C-D.** 3D renderings of the dendrites, axons, and astrocytic processes adjacent to each cilium from A and B. **E.** Every adjacent process was classified for 10 excitatory neurons and 10 astrocytes in the P36 dataset. The total processes of each type are graphed according to cell class.

To compare the type and abundance of processes outside cilia we manually identified each process that passed directly next to the cilium for 10 excitatory neuron cilia and 10 astrocyte cilia in the P36 volume. The number of adjacent axons, dendrites, and astrocytic processes for each are graphed in Figure 5E. Astrocyte processes densely populate local territories near the cell body as seen in Figure 1A and described elsewhere (Aten et al., 2022; Calì et al., 2019). Although cilia were adjacent to astrocyte processes at several points along their length, each process was counted only once. Adjacent processes from the ciliated cell were excluded, so although most astrocyte cilia had self-astrocytic processes adjacent, the value was zero. Only one of the assessed astrocyte cilia had an adjacent astrocytic process that was non-self. We found that astrocyte cilia averaged fewer adjacent axons and dendrites than excitatory neuron cilia. Because the analyzed astrocyte cilia all had ciliary pockets, we think the difference may be, at least in part, due to the reduced external cilium length. Figure S7 displays a rare observation: a cilium tip surrounded by a microglial process. Signaling molecules detected by ciliary receptors could originate in and pass through the complex extracellular environment so understanding how cilia associate with cellular processes in the neuropil might provide important clues to cilia signaling perception.

### Ultrastructure does not suggest specific interaction between dense core vesicles and cilia

The neuropeptides and neuromodulators detected by cilia have two potential sources: dense core vesicles and clear synaptic vesicles. We began by investigating whether the apparent entanglement of cilia with neuronal processes positioned cilia near dense core vesicles (DCVs). In EM DCVs contain an electron dense center that can be used to distinguish them from small clear synaptic vesicles. DCVs can be released from axons, dendrites, or synaptic termini (Pol, 2012). Occasionally DCVs appeared poised to fuse next to cilia, but typically DCVs if triggered would have released their contents near-but not directly onto - cilia. In the absence of evidence that DCVs target release directly at primary cilia we investigated whether the number of DCVs near cilia differed by cell class. To quantify and compare the number of DCVs near cilia we selected cilia in the P>270 dataset that had minimal curvature and were imaged in cross-section so that we could manually annotate DCVs within 1 μm of the center of the cilium in each image plane (for example see Figure 6A and 6B). The sample size is small because few cilia were cross-sectioned along their entire length - most turned in the imaging plane. We quantified DCVs near cilia from excitatory neurons, inhibitory neurons and astrocytes (although there is no data, we are aware of indicating astrocyte cilia contain neuropeptide receptors). We hypothesized that a longer cilium might have a higher chance of encountering DCVs so to compare the quantity of DCVs we plotted the number of DCVs as a function of length (Figure 6C). All astrocyte cilia had 10 or fewer DCVs within 1 μm, while all neuronal cilia had 10 or more. It is likely that the shielding of astrocyte cilia from neuronal processes (shown in Figure 5B and D) contributes to the reduced proximity to DCVs. In conclusion, we found that although there was little evidence for cilia-targeted release of neuropeptides, DCVs were more frequently found near neuronal cilia.

**Figure 6:**
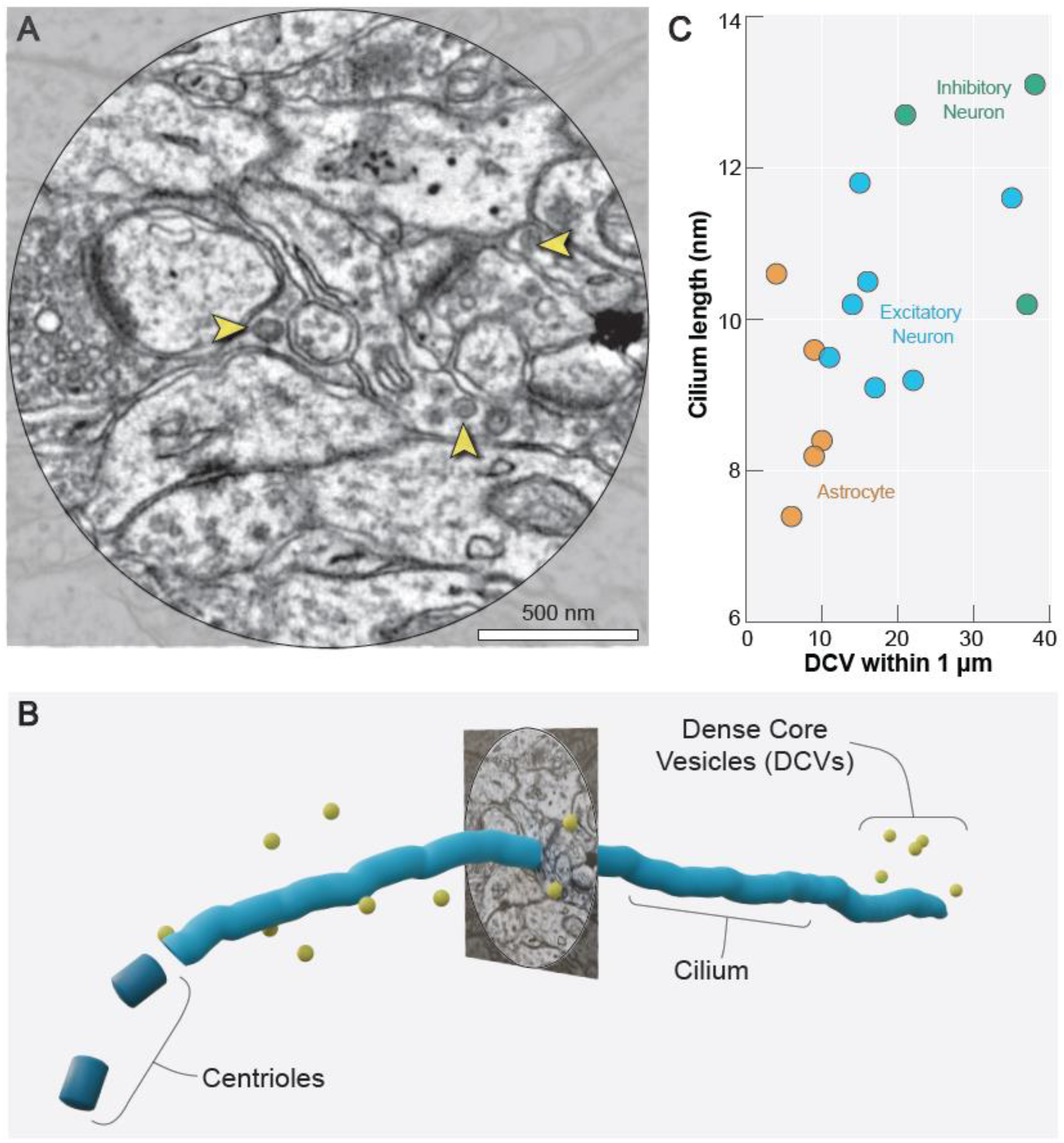
Dense core vesicles are located in the vicinity of cilia. **A.** The features within 1 μm of an excitatory neuron cilium (center) include dense core vesicles (yellow arrowheads). **B.** A 3D representation of the dense core vesicles within 1 μm of the cilium shown in A. **C.** The number of dense core vesicles within 1 μm of individual cilia is plotted against cilium length.

### Cilia are positioned to detect synaptic spillover

In addition to neuropeptides, receptors for neuromodulators that can be released from clear synaptic vesicles can also localize to neuronal cilia. To investigate the possibility that locally released neuromodulators could be detected by primary cilia, we examined synapses near primary cilia. We found synapses directly adjacent to cilia (Figure 7A upper inset and Figure S8A,B and Sup. Video 3). The ciliary membranes were seen to be as close to synapses as glia processes are and the glial processes are thought to detect spillover from the synaptic cleft. We also found synapses that were proximal to cilia, but not directly adjacent (Figure 7A lower inset and Figure S8B). In the P36 and P54 datasets, synapses have been segmented and automatically connected to their pre- and post-synaptic reconstructions (Macrina et al., 2021). Sheu et al (Sheu et al., 2022) reported that axons can synapse directly onto cilia. Although the automatic segmentation identified rare synapses onto cilia, these were determined to be false positives. Thus, we conclude that presynaptic termini all had a postsynaptic target other than the cilium (i.e., axon-dendrite synapses, axon-axon synapses, axon-soma synapses).

**Figure 7:**
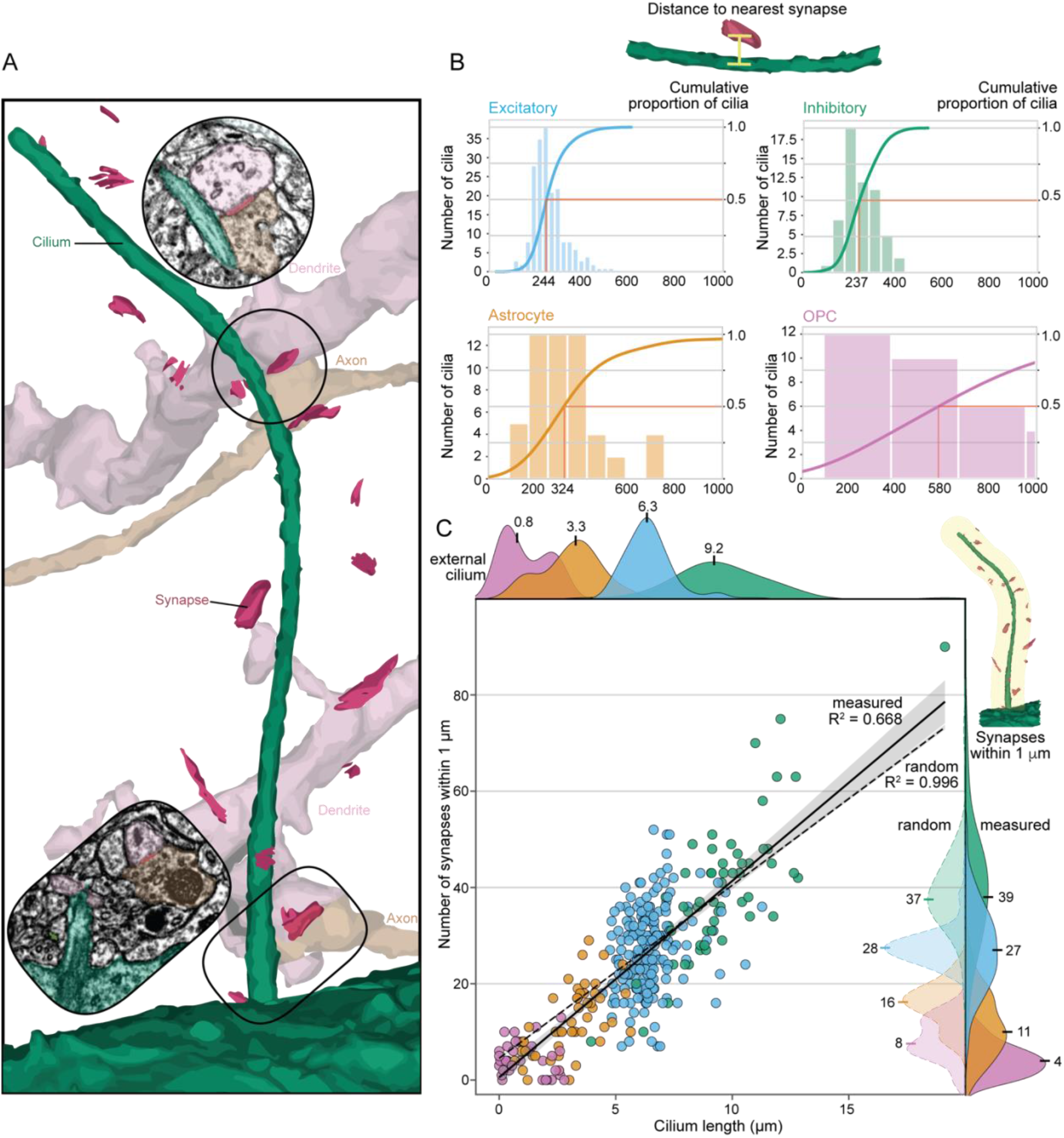
Synapses. **A.** Synapses near an inhibitory neuron cilium are shown as pink disks. The EM images to two of these synapses are shown as insets. The upper inset is a cilium directly adjacent to the cilium and the lower inset is a synapse that is proximal to the cilium. **B.** Euclidean distance to the closest synapse for each cilium in the P54 dataset by was quantified by cell class. **C.** The number of synapses within 1 μm of each P54 cilium is graphed relative to cilium length. For astrocyte and OPC cilia we plotted the number of synapses relative to the external cilium length. The solid line shows a linear regression fit across the dataset (white line, β=4.08 R^2^=0.668). Also graphed is the linear regression fit of the mean number of synapses adjacent to each cilium randomly placed in 1000 positions and orientations in the EM volume (dashed line, β=3.59 R^2^=0.996, includes the external cilia pathlength). Shading is the 95% ci. The histograms on the top show the class distributions of cilia lengths. On the right of the graph, the class distributions of the number of synapses within 1 μm are shown both as observed near cilia in the data (measured) and for comparison, as calculated if the same cilia were randomly placed in the EM volume (random).

To assess the frequency and abundance of synapses adjacent to primary cilia we used the P54 dataset, which includes the deeper layers of the cortex, to measure 1) the distance from each cilium to the nearest synapse; and 2) the number of synapses within 1 μm of each cilium. As shown in Figure 7B, half of the excitatory neuron cilia had a synapse within 244 nm. This measurement is from the center of the segmented synapse to the ciliary plasma membrane, so in many cases, the cilium may have been even closer. Inhibitory neurons were similar to excitatory neurons (median distance of 237 nm), however, the average distance to the nearest synapse from astrocyte and OPC cilia was larger (324 and 580 nm, respectively). We have already seen that astrocyte cilia have fewer adjacent processes (Figure 5) so it is possible that the ciliary pocket contributed to the increased distance to the nearest synapse.

To quantify and compare the abundance of synapses near cilia we determined the number of synapses within 1 μm of each cilium. We chose this distance because except for OPC cilia, almost all cilia had at least one synapse within 1 μm (Figure 7B). The distribution of synapses within 1 μm of each cilium is plotted for each cell class in the P56 dataset on the y-axis of Figure 7C. The inhibitory neurons had both the most synapses near individual cilia and the largest median value (39 synapses within 1 μm). Excitatory neuron cilia had a median of 27 synapses within 1 μm and astrocyte cilia and OPC cilia had substantially fewer (median values of 11 and 4 synapses within 1 μm, respectively). In addition, the number of synapses within 1 μm was also graphed relative to cilium length. Together with the pathlength of the external glia cilia, we can fit the data with a linear regression coefficient of 4.08 with an r squared value of 0.668 (Figure S8C). Using the total pathlength of glia cilia changes the coefficient value to 4.91 and r squared value to 0.606.

To determine whether synapses were enriched near primary cilia, we computationally placed each cilium mesh from the P56 dataset in 1000 random positions and orientations within the volume such that it did not collide with a cell body. We then determined the number of synapses within 1 μm and plotted the mean value for each cell (Figure S8D). The fit of the data with a linear regression coefficient was very similar to the value of the observed data (3.59 with an r squared value of 0.996). The distribution of values for each cell class, along with the fit line are included in Figure 7C. Based on this analysis we conclude that cilia were positioned to detect neuromodulators that spillover from synapses stochastically located in their proximity.

To further assess the extent to which the observed differences between cell classes were related to differences in cilia length we calculated the number of synapses per unit length of each cilium and averaged within classes. Excitatory and inhibitory neuron cilia had similar values (4.28 +- 0.19 and 4.07 +- 0.26 synapses per μm (95% ci), respectively). Astrocyte and OPC cilia had lower averages (2.42 +- 0.38 and 1.46 +- 0.36 synapses per μm (95% ci), respectively), however, when we calculated the synapse frequency for just the portion of the astrocyte that had exited the ciliary pocket, the value was similar to the neuronal cilia (4.12 +- 0.56 synapses per μm (95% ci)). Together, this analysis suggested that synapse proximity to cilia was determined by synapse abundance in the surrounding neuropil. In addition, ciliary pocket shielding accounted for the observed disparity between neuronal and glial cells. Based on this analysis we conclude that cilia are well positioned to detect neuromodulators that spillover from synapses stochastically located in their proximity.

### EM analysis reveals cell-class-specific features of cilia shape, placement, and orientation

The new knowledge that cilia can be positioned near potential signal sources suggested that the physical properties of the cilium – its length, tortuosity, position, and orientation – could influence the exposure of the cilium to external ligands. Cell class similarities in these features could suggest cellular regulation of these features. A gallery of segmented cilia meshes from P36 and P54 neurons shows similarities between cilia, while also highlighting that each is unique (Figure S9). We began by investigating how microtubule arrangements influence cilium shape. First, we examined the overall tortuosity of cilia relative to cilia length to determine whether the transition of microtubules to singlets and early terminations caused cilia to become less rigid (Figure 8A). Cilia tortuosity defined by arc-chord ratio across all cell classifications averaged 1.19 +- 0.02 (95% ci), close to the straight-line value of 1 and largely consistent across cell classifications (Figure 8A). Further, cilia pathlength did not appear to impact the gross tortuosity metric, which indicated that although the distal region of the cilium has only a few singlet microtubules (and thus a decreased persistence length) the distal region of a cilium did not become malleable. Next, we investigated whether the microtubule reductions and decreases in cilia diameter shown in Figure 4 B-D altered cilium shape. We plotted the binned local curvature, defined as the inverse radius of a circle tangent to the skeleton’s path, along the length of each cilium by cell class. On the same axis, we plotted the cilium diameter (Figure 8B). The value of local curvature increased at the base of the cilium as the diameter decreased in cilia from both classes of neurons and in astrocyte (Figure 8B). Both the diameter and the local curvature leveled out at a similar distance from the base of the cilium. While OPC cilia diameter drops off very close to the base, the other trends in diameter stability and curvature leveling are not apparent (which could in part be related to the internal cilia vesicles shown in Figure S2. Taken together, these data indicate that the changes in A-tubule staining, and the B-tubule termination in the proximal cilium facilitated cilium bending. The rate of local curvature appears sustained in the distal cilium preventing longer cilia from becoming increasingly tortuous.

**Figure 8:**
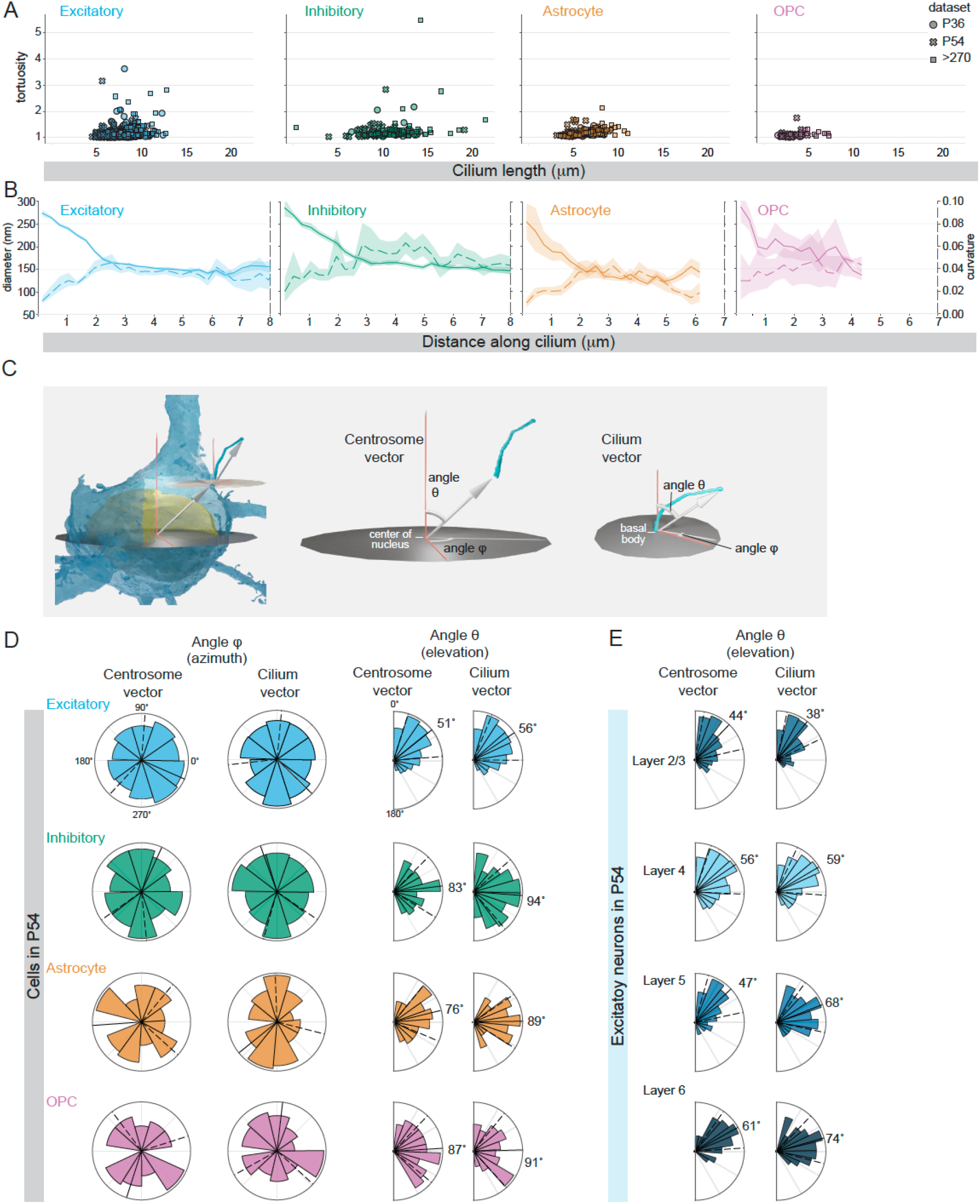
Cilium shape, placement, and orientation can be stereotyped within a cell class. **A.** For each cilium analyzed, the tortuosity is plotted relative to cilium length. **B.** The diameter along the length of each cilium in the P54 dataset are plotted on the left y-axis (solid line) and the local curvature of each cilium is plotted using the right y-axis (dotted line). For each the 95% confidence interval is indicated by the shading around the line. The value of the diameters within the first 5 μm are also presented in Figure 6. **C.** Two vectors were created to analyze cilium placement and orientation: the centrosome vector originates at the center of the nucleus and extends to the mother centriole while the cilium vector extends from the base of the cilium to the tip. The azimuth is represented by the angle **φ** and the elevation by the angle **θ**. The spherical coordinate system is defined by a zenith representing an axis from white matter to pia. **D.** Polar plots show the distribution of centrosome and cilia vectors for each cell in each class in the P54 dataset. The value of the circular mean is indicated and represented by a solid black line and the dotted lines represent the circular standard deviation around the mean. **E.** The centrosome and cilium vectors of Excitatory neurons in each layer are plotted.

Finally, we sought to determine if there were any cell class similarities in centriole placement and cilium orientation. Immunofluorescent imaging of cilia in the hippocampus and cortex has provided evidence that cilia within populations of neurons can align. Cilia in CA1 have been shown to orient along the apical-basal axis (either base-to-tip or tip-to-base) (Sheu et al., 2022) and in the cortex cilia orient toward the pial surface (Kirschen et al., 2017). To investigate possible similarities within or between cell classes in centriole positioning or cilia orientation we used two vectors: 1) from the center of the nucleus to the mother centriole that we call the centrosome vector, which defines where on the cell the cilium originates; and 2) a vector that extends from the base of the cilium to the tip that we call the cilium vector, which defines cilium orientation (Figure 8C). For each vector we plotted the angle phi (a measure of the direction of the vector angle relative to the rostral-caudal and anterior-posterior axis) and the angle theta (a measure of elevation or depression that indicates if the vector is pointing toward the pia or toward the deeper layers of the tissue, i.e., dorsal or ventral). The distribution of the angles for each cell class in the P54 dataset is presented in Figure 8D. We observed no uniformity in the angle phi; phi was distributed throughout the possible angles for all cell classes. In contrast, for both the centrosome and cilia vectors of excitatory neurons the angle theta condensed around an average of 51° and 56°, respectively. In the physiological context, the polarized centriole vector meant that the mother centrioles docked near the base of the apical dendrite. The polarized cilia vector meant that on average, cilia tended to orient toward the pia. Because the P54 dataset extends through Layer 6, we were able to analyze and compare the cilia and centrosome vectors between cells of the same class at different depths. The distribution of values for the excitatory neurons in each layer are plotted in Figure 8E. We found that the positioning of the cilium origin and the directionality of the cilium vector shifts away from the zenith across the visual cortex (for example, the cilium vector shifts from 38° in layer 2/3 to 74° in layer 6). Taken together the analysis of the physical properties of the cilium reveal that centrosomes and microtubules (interior structures) influence the shape and trajectory of the cilium, which could in turn influence the extracellular environment surrounding the cilium.

## Discussion

Volumetric EM datasets made it possible for us to locate and examine hundreds of cilia in an array of orientations that would have been previously unimaginable. In conventional studies, distal cilia with singlet microtubules and narrow diameters would not have been identified as cilia because they deviated from the stereotyped 9 + 0 doublet organization. These new views revealed cell-type specific differences in cilia ultrastructure. In addition, we have had the unprecedented opportunity to explore the 3D environment outside cilia – the context for signal transmission and detection. Together with quantitative analysis of primary cilia position and orientation, we have a new understanding of how cell-specific structural features could define cilia exposure to external signals.

The analyzed datasets were generated with 4 nm x,y resolution, which is sufficient to resolve synaptic connections, although EM is capable of sub-nanometer resolution. However, because TEM is a non-destructive method and many labs generating large TEM datasets store and label their sections to provide a kind of physical archive, we were able to locate and reimage the same structures at higher resolution. This is the first example we are aware of where someone has been able to enhance a large processed and reconstructed volume EM dataset by introducing higher-resolution data to address a new question. It was also important that we were able to examine cilia in EM volumes from three animals. While not replicates, the similarities we observed gave us confidence while the differences between datasets indicated variability, possibly attributable to age and/or cell layer differences, which can be further explored. When we began annotating cilia in the unsegmented P>270 dataset, it was impractical to try to identify the many processes surrounding cilia in the neuropil. The 3D cell membrane and synaptic cleft segmentations of the P36 and P54 datasets gave us the ability to examine and identify processes using information tens or hundreds of microns away. Instead of being limited to measurements of cilia pathlength, we assigned 3D volumes representing the shape of each cilium which enabled us to measure cilia diameter and the distances between the cilium surface and external objects. Specifically, 3D databasing and computational tools based on synaptic cleft segmentation allowed us to collect metrics about the relative placement of synapses to the external geometry of cilia. Larger and more diverse 3D EM datasets are now being produced across species and regions. In the future, the annotations in this study may serve as ground truth data to train networks to locate cilia across the brain. Here, the detailed examination and quantitative assessment of individual cilia revealed unanticipated cilia diversity in the visual cortex.

We previously had little idea how cilia in the visual cortex were anatomically distinct. We found that cilia differed in length, origin structure, TZ length, microtubule organization, and the presence of internal cilia vesicles. The division between cells with pocket cilia versus surface cilia suggests different ciliogenesis pathways are employed in different cells. These features may be key indicators of cilia biogenesis and function that can be evaluated in many contexts. For example, the sub-population of astrocytes that had surface cilia instead of pocket cilia could be functionally distinct. Neurons outside the visual cortex may use the internal ciliogenesis pathway (for example a Purkinje neuron has a pocket cilium (Cerro et al., 1969). In the mouse optic nerve, all the astrocytes examined by serial block face scanning EM had concealed or pocket cilia and OPC cilia were docked at the surface (Ono et al., 2022). The observed differences between neuron and astrocyte transition zones may also deviate in other brain regions and transition zones might even change over time as seen in other systems (Akella et al., 2019; Wiegering et al., 2018). It is also possible transition zones could be more variant in other brain regions. In *Drosophila,* the immotile cilia of olfactory neurons and the motile cilia in auditory neurons have different Y-link structures (Jana et al., 2018). Additional studies will be needed to determine if the measurable differences between neuron and astrocyte base structures and TZs contribute to functional specialization. We also speculate that previously unrecognized structural differences between cell types could explain why mutation of genes coding for cilia and centrosome proteins can have diverse outcomes.

The differences in microtubule structure both provided new insights and raised many questions. The arrangement of microtubules differed between neuronal and astrocyte cilia. A possible explanation for this is that the cell classes express different tubulin monomers (Silva et al., 2017). For example, beta tubulin 3 is often used as a neuron marker because it is absent from glia. Perhaps different complements of tubulin monomers create structurally distinct cilia. We also discovered densities present in the A-tubule near the base of cilia. Experiments are needed to determine if, as we hypothesize, the densities represented microtubule inner binding and whether their presence prevents B-tubule termination. The observed configurations of doublet and singlet microtubules also revealed gaps in our knowledge about IFT movement, which has been shown to be tubule-dependent in motile cilia (anterograde transport on the B-tubule and retrograde transport on the A-tubule) (Stepanek and Pigino, 2016). In *C. elegans*, alternate motor configurations conduct anterograde transport on distal microtubule singlets (Snow et al., 2004). The issue of IFT in the absence of B-tubules, has been raised by (Sun et al., 2019) who describe microtubule doublet to singlet transitions as a part of a 3D model of cilia microtubules from serial section electron tomography of cultured epithelial cell cilia. In that study, they also predict that densities bridging microtubules to each other or to the ciliary membrane are microtubule associated proteins (MAPs). Identification and characterization of MAPs in cilia will be necessary to determine the extent to which each supports or undermines microtubule stability. How microtubules, microtubule inner binding proteins, and MAPS together create cilia stability may be dynamic as evidence suggests cilia can change through development and through a single day.

While the views inside cilia provided important insights into cell-class specific differences in cilia structures, the quantification of structures outside cilia suggested novel parameters with potential to influence cilia signaling. The absence of such evidence that targeted secretion of DCVs occurs near primary cilia is consistent with the prevailing paradigm that peptide signaling occurs largely through bulk flow in the neuropil. Contents of DCVs are considered long-lived in comparison to contents released from clear synaptic vesicles which are often cleared by machinery within the synapse or by proximal astrocytic processes that also detect spill-over. Cilia adjacent to synapses appear to have as much access to spill-over as an adjacent astrocytic process would. Because receptors for both dopamine and serotonin can be localized to cilia and both are synapse-released neuromodulators, we hypothesize that when appropriate receptors are trafficked to cilia, signaling events at synapses can be detected by primary cilia. Because synapses were not enriched around cilia, we hypothesize that cilia are sensing general synaptic activity. This model is different from the proposed detection of serotonin by cilia in the hippocampus where axons were reported to synapse directly onto cilia. In the visual cortex we did not observe any synapses directly onto cilia; all presynaptic termini near primary cilia had a separate post-synaptic target. It is possible that brain-region differences account for the different observations. These investigations have given us new understanding of the nature of the environment surrounding cilia. Some features, like ciliary pockets or glial processes might shield cilia from potential ligands, while others, especially cilia adjacent synapses, could be sources for a subset of cilia signals.

We are just beginning to understand how primary cilia influence neuronal and glial activity (Tereshko et al., 2021; Wang et al., 2021b). A recent study examining primary cilia in neurons and astrocytes in an EM volume of the human anterior temporal lobe supports many of our conclusions (Wu et al., 2023). Along with other recent studies (Kiesel et al., 2020; Ono et al., 2022), the ultrastructural analysis presented here demonstrates that primary cilia structures cannot be stereotyped. Furthermore, although synapse abundance near cilia may be stochastic, in combination with placement of the cilium origin, regulation of cilia structural features can determine the extracellular environment monitored by cilia localized receptors.

## Acknowledgements

C.M.O., D.B. and J.L.S. were supported by the Howard Hughes Medical Institute. N. M. C. acknowledges support from NSF NeuroNex 2 award 2014862. W.C.A.L. acknowledges support from the NIH (MH117808). We thank Tom Kazimiers from kazmos GmbH for supporting our work by contributing open-source code to CATMAID and managing the instance used for annotation. We are also grateful to Jennifer Garrison and the Les Treilles Foundation for bringing collaborators together. We also thank Christina Gladkova, Lauren Porter, Andy Moore, Chris Obara, Casey Schneider-Mizell, and Kayla Jewett for helpful discussions and comments on the manuscript. We thank the Allen Institute for Brain Science founder, Paul G. Allen, for his vision, encouragement, and support.

## Author contributions

C.M.O., R.T., N. C. and J.L.S. conceived the project and wrote the paper with inputs from all authors. C.M.O. annotated and analyzed EM data. C.M.O. and R.T. analyzed the data. T.S.K. located, and re-imaged sections with help from W.C.L and A.K. who also processed and aligned images. D.B. helped with early annotation of the P>270 dataset and J.B., L.E., S.S., A.L.B., F.C., D.B., C.M.O., and N.C. provided cell identification and classification for the P36 and P54 datasets.

## Methods

### Overview of TEM datasets

The TEM volumes used in this study were collected as a part of collaborative projects designed to determine neuron connectivity. The P36 and P>270 mice have been described (Schneider-Mizell et al., 2021; Bock et al., 2011; Dorkenwald et al., 2022). The P54 mouse was a cross of B6;CBA-Tg(Camk2a-tTA)1Mmay/J (Jax: 003010) and B6;DBA-Tg(tetO-GCaMP6s)2Niell/J (Jax: 024742). The age of this mouse at perfusion was P54. All animal procedures were approved by the Institutional Animal Care and Use Committee at the Allen Institute for Brain Science or Baylor College of Medicine. As detailed with the publication of the original dataset (Bock et al., 2011) i*n vivo* fluorescent calcium imaging preceded isolation of a volume of the visual cortex from mice. The tissues were then stained, sectioned, and imaged with 4 nm resolution in XY. Section thickness, and thus z resolution is 40 nm. The tissue volumes were computationally reconstructed by stitching and aligning these images to create a large volume. Neurons were then traced as skeletons directly in the volume (Bock et al., 2011), or densely segmented and proofread to generate 3D reconstructions (Turner et al., 2022). The heavy metal staining and resolution that makes electron microscopy suitable for neuronal and synaptic reconstruction is sufficient to resolve many intracellular features including nuclear pores, mitochondria cristae, polysomes and microtubules, as well as synaptic vesicles and pre- and post-synaptic densities. Distinguishing features of each dataset including each animal’s sex and the area of visual cortex imaged are listed in Supplemental Table 1.

### Cell Classifications

Cell features provided the information necessary for cell classification as described in (Peters et al., 1991). Excitatory pyramidal neurons were distinguished from the inhibitory neurons based on characteristics including nuclear morphology, axon and dendrite positioning and morphology and the post-synaptic density adjacent to axon termini. In EM, Astrocytes were identified by the dark ring of chromatin around the periphery of the nucleus, the presence of glycogen granules and the many processes that appear to seep rather than project into the surrounding spaces. In segmented data astrocytes were also identified by the characteristic overall cell morphology. Microglia and oligodendrocytes both have small dark nuclei and dark staining cytoplasm but differ in the diameter and content of the processes: oligodendrocyte processes are thin and packed with microtubules while microglia processes narrow more gradually. OPC cells have lighter cytoplasmic staining than oligodendrocytes and microglia. For more details about the identification and characteristics of OPC cells see (Buchanan et al., 2022). In P>270, Astrocytes in P>270 were classified based on the presence or absence of abundant dense intermediate filaments as protoplasmic or fibrous respectively. Many fibrous astrocytes were in cortical layer 1. Within the P54 dataset every cell was assigned a cell classification and layer initially based on a clustering analysis of nucleus features (Elabbady et al., 2022). We reconstructed cells based on this initialization then manually confirmed classifications using the morphology of segmented processes. For analysis, layer boundaries in the P54 dataset were defined using a set of manual point annotations based on the size and density of pyramidal cells for each layer to generate average planes dividing L1, L2/3, L4, L5, and L6. Although the P54 volume encompasses layers 1-6, layer 1 was excluded from the 3D segmentation, and thus from our analysis.

### Annotation of cilia and centrosomes

For the P36 and P54 volumes, the data was accessed through Neuroglancer (neuroglancer) and point annotations were used to mark the cilia base and tip, centrosome position, cilium exit from pocket cilia and vesicles. Initial annotation and segmentation of P>270 was done using Trak EM2 (Cardona et al., 2012). Annotations were migrated to CATMAID (Schneider-Mizell et al., 2016; Saalfeld et al., 2009) and all annotating continued in CATMAID. To facilitate work across multiple datasets, we contracted kazmos GmbH to extend CATMAID to access and annotate data in neuroglancer precomputed format. This allows all datasets released in this format, including many new and upcoming large EM volumes, to be accessible and annotated from local CATMAID instances without the need to copy data locally. We used CATMAID to trace ciliary skeletons from base to tip. To locate cilia, we used either the 3D cellular models to identify cilia and then navigated to the corresponding EM images or we searched the somata in the EM volume for centrosomes and then looked to see if cilia were present. Incomplete cilia were included for measurements of cilia frequency but excluded from other analysis. P36 is the only volume in which every cell in the volume was assessed. In P54 and P>270, cilia in a subset of the abundant excitatory neurons and glia were annotated and all or almost all inhibitory neurons were annotated.

There were astrocytes and neurons in which we were unable to locate a cilium, but these cases were very rare (see table 1). Cells that left the volume or that had large imaging gaps were excluded from analysis if a ciliated centriole was not found. In the two nonciliated astrocytes in P36 we found both centrioles in the cytoplasm and are confident that these are nonciliated cells. We are less confident about the absence of cilia in the inhibitory neurons that are not included in the ciliated fraction because we found no centrioles in these cells. It is possible that ciliated centrioles are present and were either overlooked (despite multiple efforts) or obfuscated by imaging gaps or artifacts. In all microglia we found two centrioles. In P36 and P54 all the oligodendrocytes had two nonciliated centrioles, however, no centrioles were found in any of the oligodendrocytes in P>270. We are not aware of any reports of centriole loss in oligodendrocytes and wonder if the difference could be due to the age difference in the mice. Rare neurons with multiple primary cilia have been found in the hypothalamus (Koemeter-Cox et al., 2014). In V1 we observed only monociliated neurons. We did find an astrocyte with cilia extending from both centrioles and another ciliated astrocyte that had a large vesicle associated with the second centriole that appeared to be a pre-ciliary structure. Endothelial cells, part of the brain circulatory system, were omitted from our analysis, however, we did locate pairs of nonciliated centrioles in 3 pericytes in the P>270 dataset.

### Manual Segmentation in Amira

Cilia, ciliary vesicles and the cytoplasm were manually segmented using Amira 3D version 2021.1 (Thermo Fisher Scientific). Cropped image stacks were imported. Next, the cilia structures and cytoplasm were each selected using the lasso tool in Amira in individual layers. were created by manually tracing structures. The resulting segmentation was converted into a triangular mesh surface using the ‘Generate Surface’ module in Amira with constrained smoothing and a smoothing extent of 9.

### Re-imaging TEM sections at higher resolution

TEM has the advantage of improved microtubule contrast and the ability to revisit a sample and image again at higher magnification, which is not possible with destructive volume EM methods, such as serial block face scanning EM (SEM) and FIB-SEM (focused ion beam-SEM). Furthermore, TEM enables higher resolution imaging than SEM methods. To obtain higher quality images of cilia identified in the P>270 dataset (Bock et al., 2011), we reimaged the same samples with a TEM at higher magnification (JEOL 1200EX). The samples consisting of 1,215 serial sections on Pioloform-coated TEM slot grids were stored in a nitrogen dry box for more than 10 years since the initial imaging. Yet, the new images were comparable to the originals, suggesting that there was minimal sample deterioration during this storage period. Using the published image volume and the original imaging notes as a guide, we identified which grids contained the cilia of interest and located them within each grid. The cilia were imaged at much higher magnifications (30,000x to 65,000x) than previously. Since cilia can extend several microns in length, some cilium required re-imaging from up to 100 sections. For each cilium, the high-resolution serial images were then aligned together using an elastic alignment software pipeline (AlignTK).

### Transition zone measurements

Cilia in the P>270 dataset that were sectioned close to parallel to the imaging plane at the cilium base were identified. To measure transition zones, the density between the ciliary membrane and the axoneme was measured in every section where it was visible. Because the transition zone is a periodic structure, the density was not visible in every plane. The variability in measurements from an individual cilium is likely because the cut angle was not precisely parallel.

### Visualization of 3D objects using Blender 3.1.2

3D meshworks of cells or processes were imported as wavefront OBJ files. Where necessary, cilia meshworks were isolated and colored separately. For glial cells, the portions of the cell obstructing the view of the cilium were cut away.

### Quantification of processes adjacent to cilia

Using Neuroglancer it was possible to navigate to the 2D view anywhere along each 3D structure. Astrocytic processes were identified based on their characteristic morphology and the presence of glycogen granules. Axons were followed away from cilia until 2 or 3 presynaptic termini were identified. Similarly, the identity of dendrites, which typically have characteristic spines, were verified by locating postsynaptic densities. For 10 astrocytes and 10 excitatory neurons, every process adjacent to the cilium was classified, colored according to type and quantified.

### Quantification of dense core vesicles near cilia

Cilia in P>270 that were sectioned perpendicular to the imaging plane along the entire cilium were identified. A cylinder with a 1 μm diameter was created around the annotated cilium pathlength. Each DCV within the cylinder was annotated and the total for each cilium recorded.

### Skeleton generation and processing

Skeletons for P36 and P54 were generated using a semi-automated iterative process based on point annotations of the tip and base as well as the mesh derived from the neuronal segmentation. These skeletons were generated by taking a bounding box cutout of a mesh and filtering it by the distance from a line segment between the base and tip of the cilium, then skeletonizing the largest resulting connected component. Subsequent steps filtered the mesh based on a reduced distance from the mesh to the last computed skeleton. Skeletons at each step were generated by thinning with mesh contraction as implemented by the skeletor package (Schlegel et al., 2022) followed by mesh-based TEASAR and ray-tracing based diameter estimation (Dorkenwald et al., 2020). The resulting skeletons were automatically checked for connectivity to manually defined base, tip, and exit points and reviewed in CATMAID. This process results in a skeleton that approximates the medial axis of the cilium as well as a representative compartment submesh. Skeletons in the P>270 were generated manually by placing nodes at the center of the cilium in each plane.

### Analysis of annotated data

Analysis of multiple datasets required methods for integrating results across different annotation and data sharing platforms including reading skeletons, meshes, and metadata from a variety of data sources. A python package “pycilium” (github.com/AllenInstitute/pycilium) developed for this project is publicly available on github that allows retrieval of annotations generated using CATMAID, trakEM2, and pre-defined neuroglancer state schemas and supports saving versioned copies of data relevant for subsequent analysis.

### Orientation calculations

Ensuring a consistent coordinate space for comparing EM datasets was a non-trivial problem. To address the fact that the V1 datasets used in this study were not natively rendered in a consistent orientation, each dataset was assigned a 3D rigid transformation defined to align the mean orientation of manually defined axon initial segments in layer 2/3 to a consistent rectilinear axis across datasets. This axis, directionally representing the depth from pia to white matter, was used as the zenith when defining spherical coordinate systems and as the axis along which depth was calculated.

Directional mean and directional standard deviations were found using the following formulations:

Means of trigonometric functions were calculated by averaging the sin and cosine of angles:

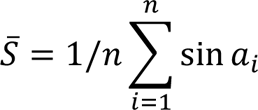

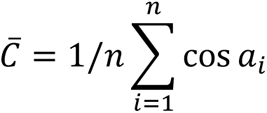

Angular mean was described using the two-input arctangent of the sin and cosine means

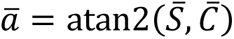

Angular standard deviation was found with the following formulation:

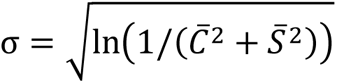

### Morphological calculations

Prior to morphological calculations skeletons were smoothed using a gaussian kernel to mitigate the effects of jitter that arises from inconsistencies in reconstruction and dataset alignment. Tortuosity was calculated as a gross metric ratio of skeleton path length to vector sum length, resulting in possible values [1, inf). Curvature was defined as an instantaneous value describing the inverse of the radius of curvature along a curve. This calculation was based on a standard formulation defined by the following equation, where gamma is the arc-length parameterization for the curve, implemented in python using the gradient of the smoothed skeleton.

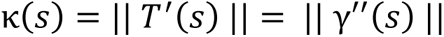

### Synapse data analysis

The P36 and P54 datasets are linked to volumetric segmentations representing the synaptic clefts of all synapses in each volume. To interrogate the spatial relationship between synapses and cilia in neuropil, we queried the 3D skeleton representations of cilia against a postgresql database table with postGIS support schematized to represent the 3D extents and center points of the segmented synaptic clefts. These queries properly scaled between anisotropic and down sampled segmentation spaces to quantify the distance and distribution of synapses near cilia. An R-tree representation of synaptic and somatic bounding boxes was used to simulate the effect of placing different model cilia randomly in the volume. Each model cilium, derived from a proofread full cilium from the dataset, was subjected to a rigid transformation defined by a 3D translation picked from a uniform distribution across the bounding box of the synaptic segmentation as well as a random 3D rotation defined by the QR decomposition of a uniformly random 3D matrix. Model cilia were excluded and the transformation recalculated if they fell within the meshes defining the somatic compartment.

