## Supplemental figures and legends for "Nanometer-scale views of visual cortex reveal anatomical features of primary cilia poised to detect synaptic spillover"

### Supplemental Figures Legends

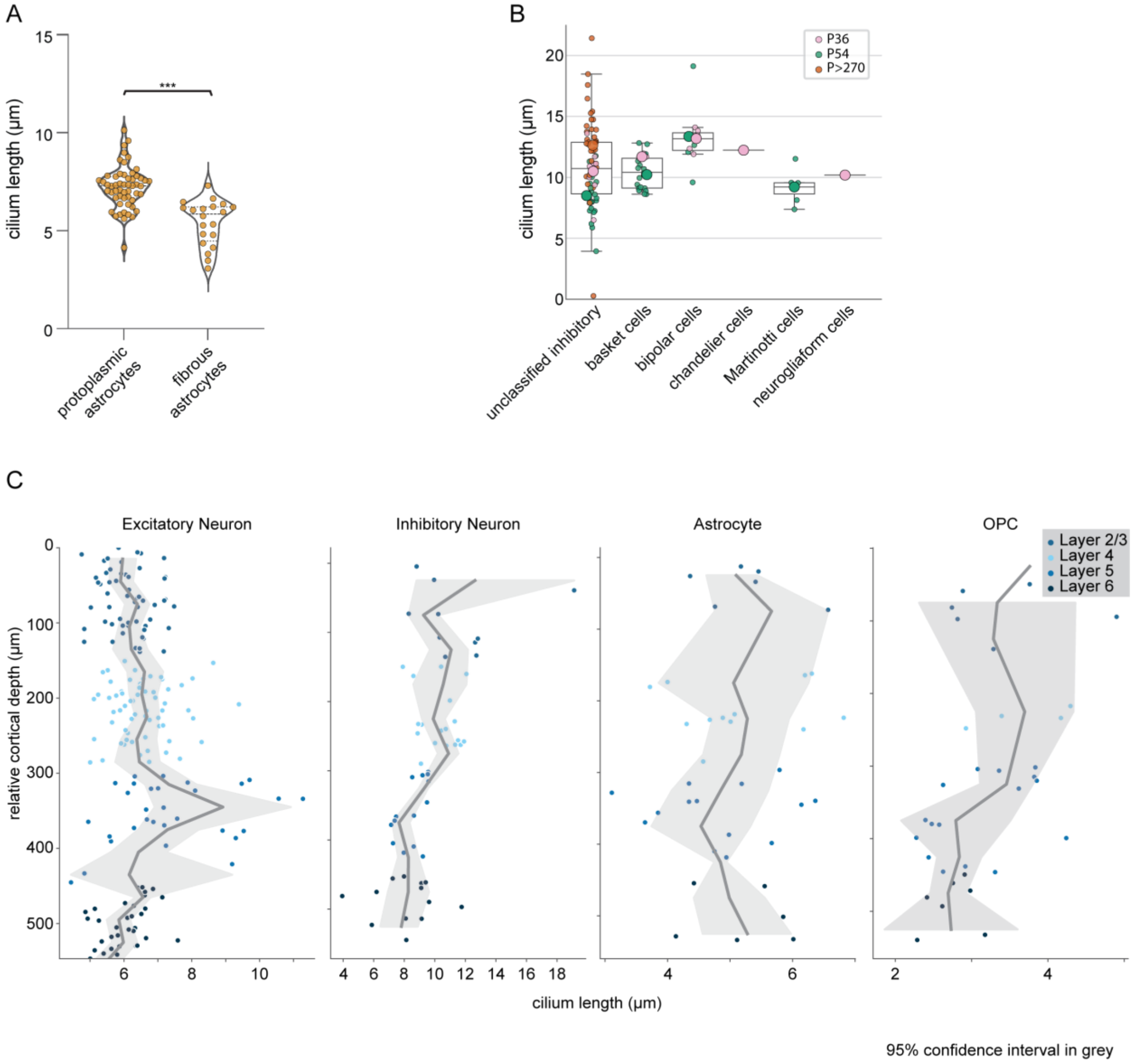


**Supplemental Figure 1: Cilium lengths differ between cell types within cell classes**

**A.** Fibrous astrocytes in the P>270 dataset were distinguished from protoplasmic astrocytes based on the presence of abundant dense intermediate filaments. Measured cilia lengths for each are plotted. P value = 0.0001. **B.** Inhibitory cell type identities were available for several inhibitory neurons in the P36 and P54 datasets. Cilia lengths of all unclassified inhibitory neurons are compared to the known cell types. The circle color indicates the dataset of each neuron. The large circles superimposed are the average value from the measurements from each animal. **C.** Cilia length for each cilium in the P54 volume is plotted relative to cilium depth by cell class. The cilium layer is indicated by color. The grey shading indicates the 95% confidence interval for the trendline.

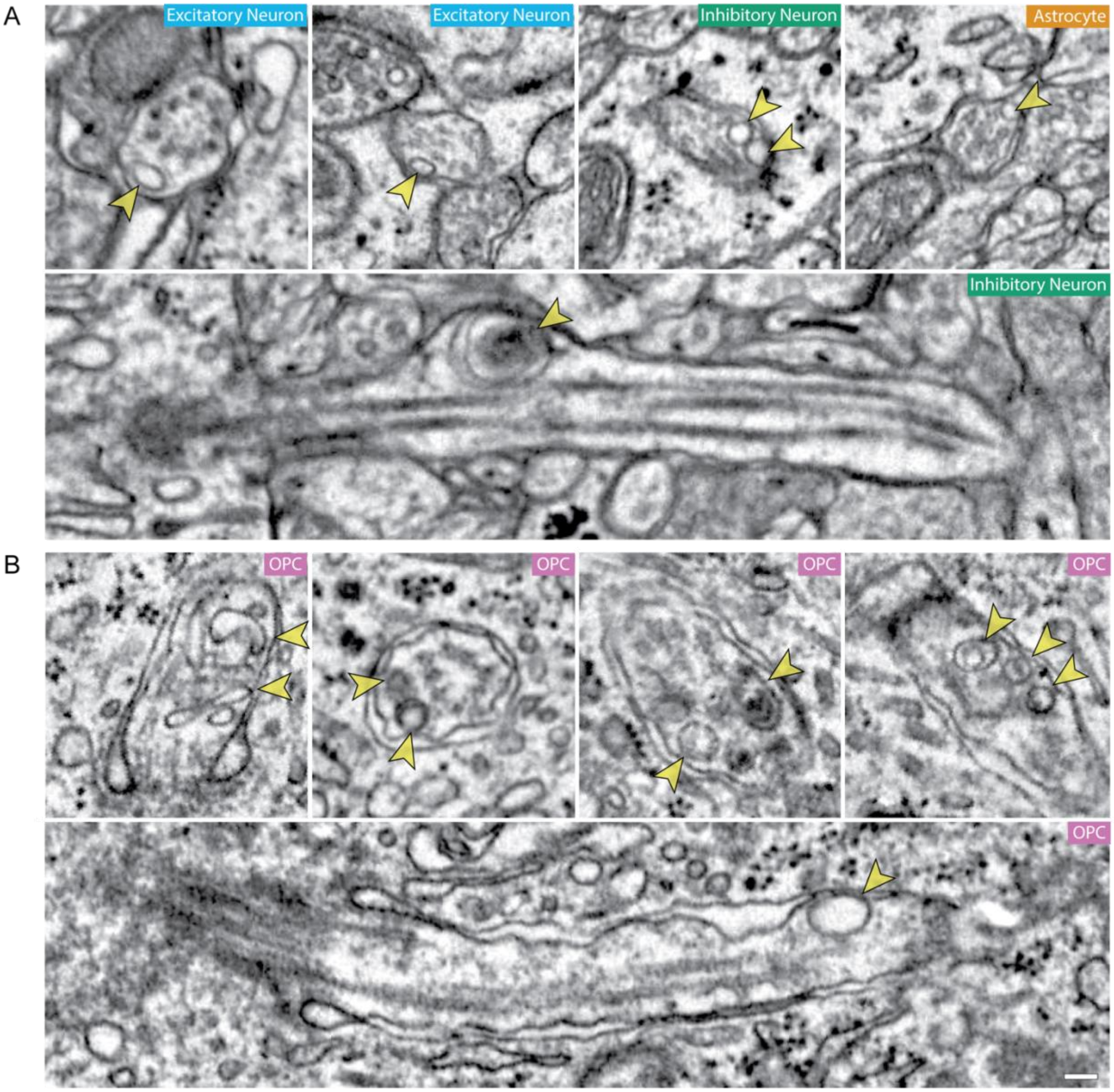


**Supplemental Figure 2: Internal cilia vesicles, occasionally found across all cell classes, were enriched in OPC cilia.**

**A.** Vesicles were occasionally discovered within both neuron and astrocyte cilia (yellow arrowhead). **B.** OPC cilia contained diverse membrane structures including single and double membrane structures with translucent and electron dense centers. Scale bar is 100 nm. Scale bar is 100 nm.

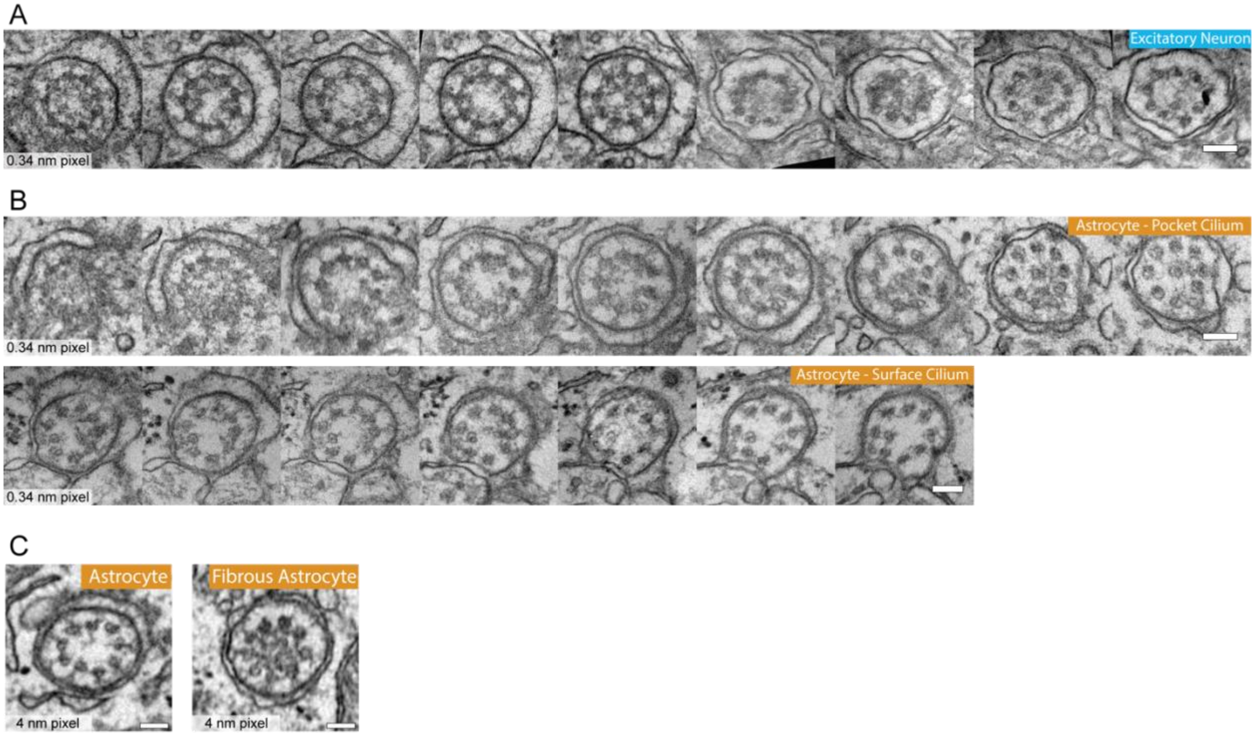


**Supplemental Figure 3: High resolution views of neuron and astrocyte cilia transition zone cross-sections**

**A and B.** Z series progressions of the transition zone at the base of an excitatory neuron (A) or astrocytes (B). **C.** Cross-section views of astrocyte transition zones. Note the presence of electron staining in the center of the fibrous astrocyte cilium. Scale bars are 100 nm.

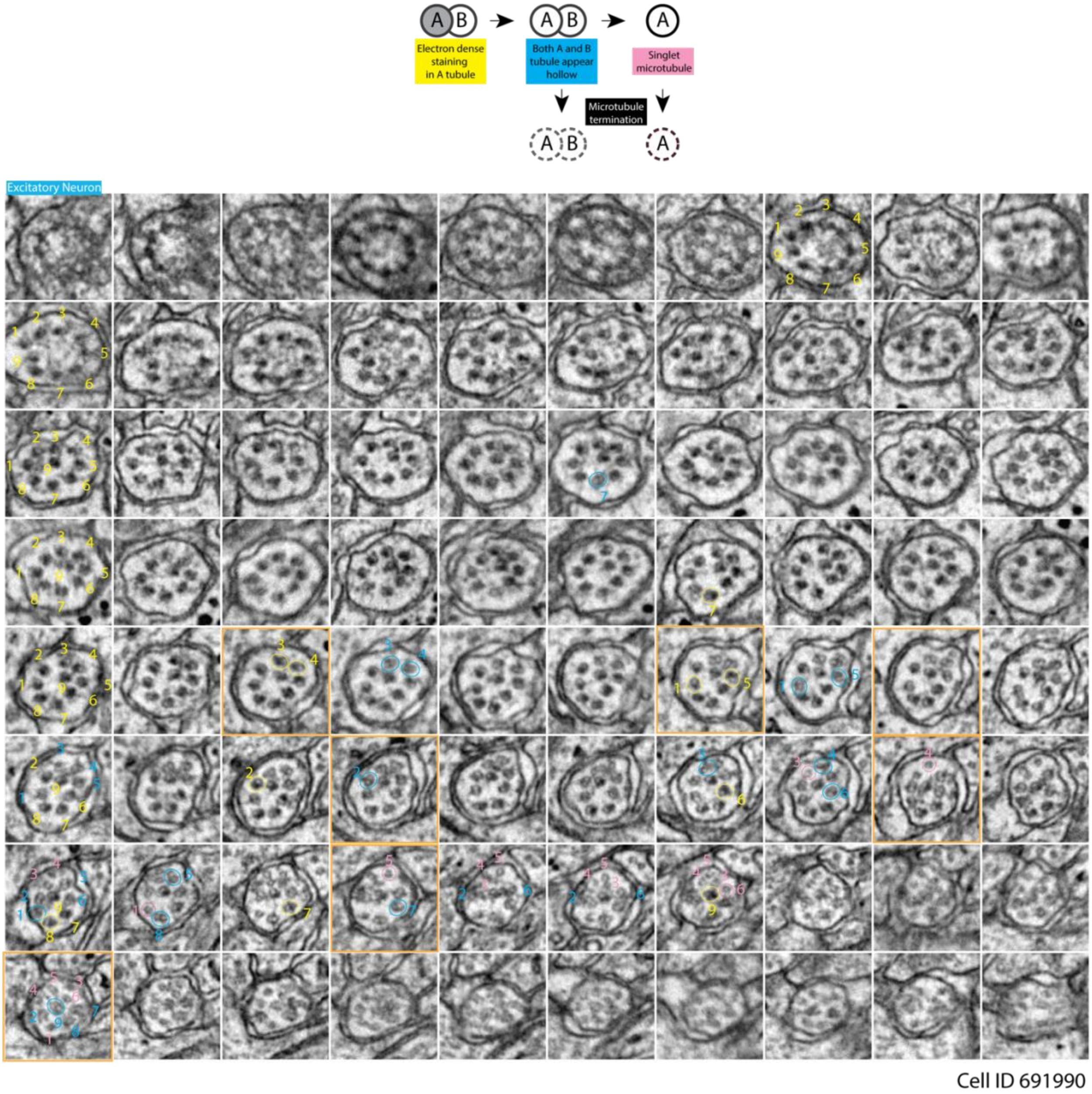


**Supplemental Figure 4: Microtubule changes at the base of an excitatory neuron cilium**

Serial sections of the excitatory neuron shown in Figure 4B, and Sup Video 2 are arranged in rows. Microtubules doublets are numbered in yellow to indicate that these are doublets that have electron-dense staining in the A-tubule. Blue circles indicate the transition to microtubules with a translucent B-tubule and pink circles indicate where singlet microtubules become clearly visible. The panels shown in Figure 4B are outlined in orange. Each panel is 400 x 400 nm.

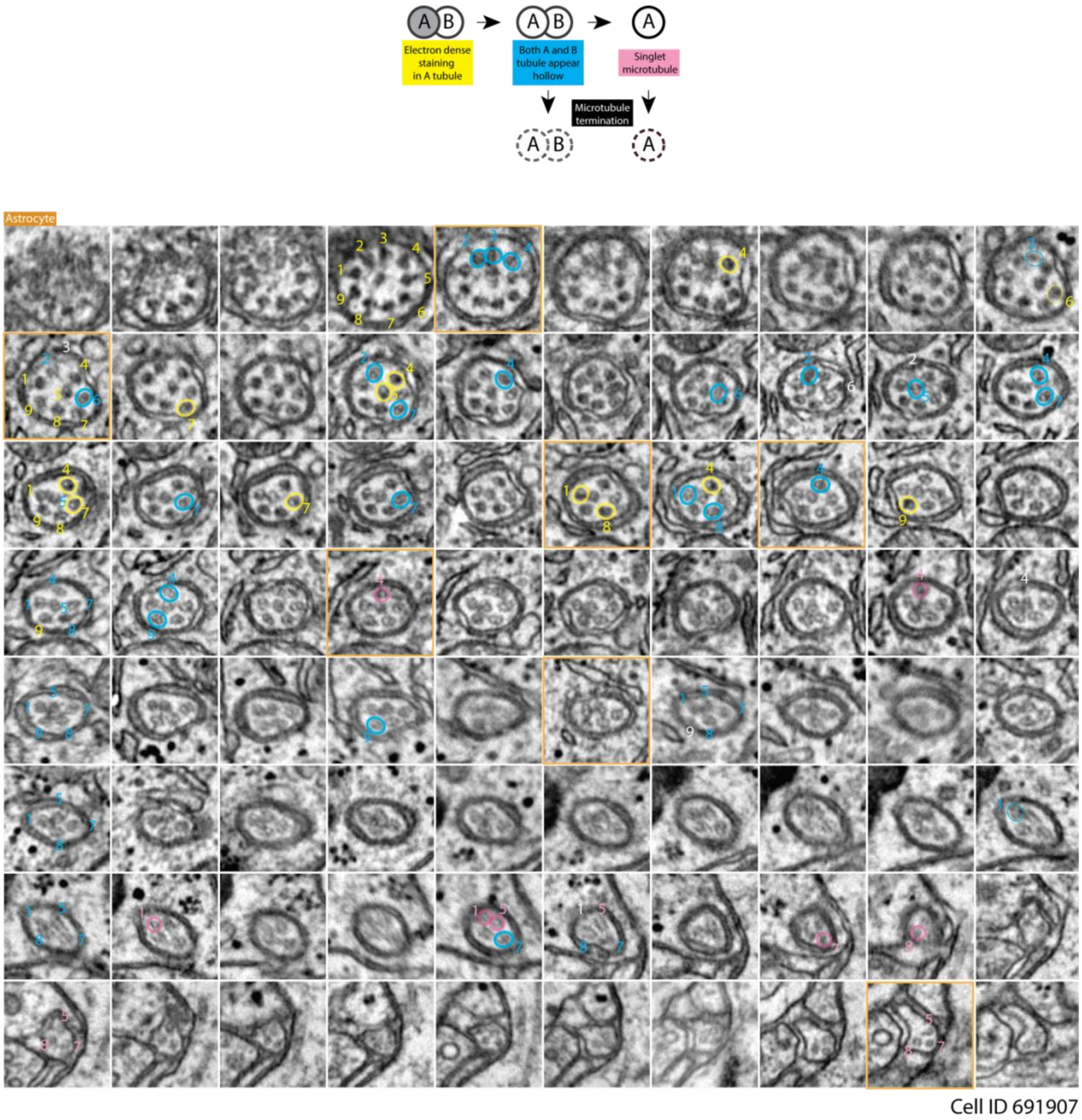


**Supplemental Figure 5: Microtubule changes at the base of an astrocyte cilium**

Serial sections of the excitatory neuron shown in Figure 4C, and Sup Video 2 are arranged in rows. Microtubules doublets are numbered in yellow to indicate that these are doublets that have electron-dense staining in the A-tubule. Blue circles indicate the transition to microtubules with a translucent B-tubule; pink circles indicate where singlet microtubules become clearly visible; and white numbers indicate microtubule termination. The panels shown in Figure 4C are outlined in orange. Each panel is 400 x 400 nm.

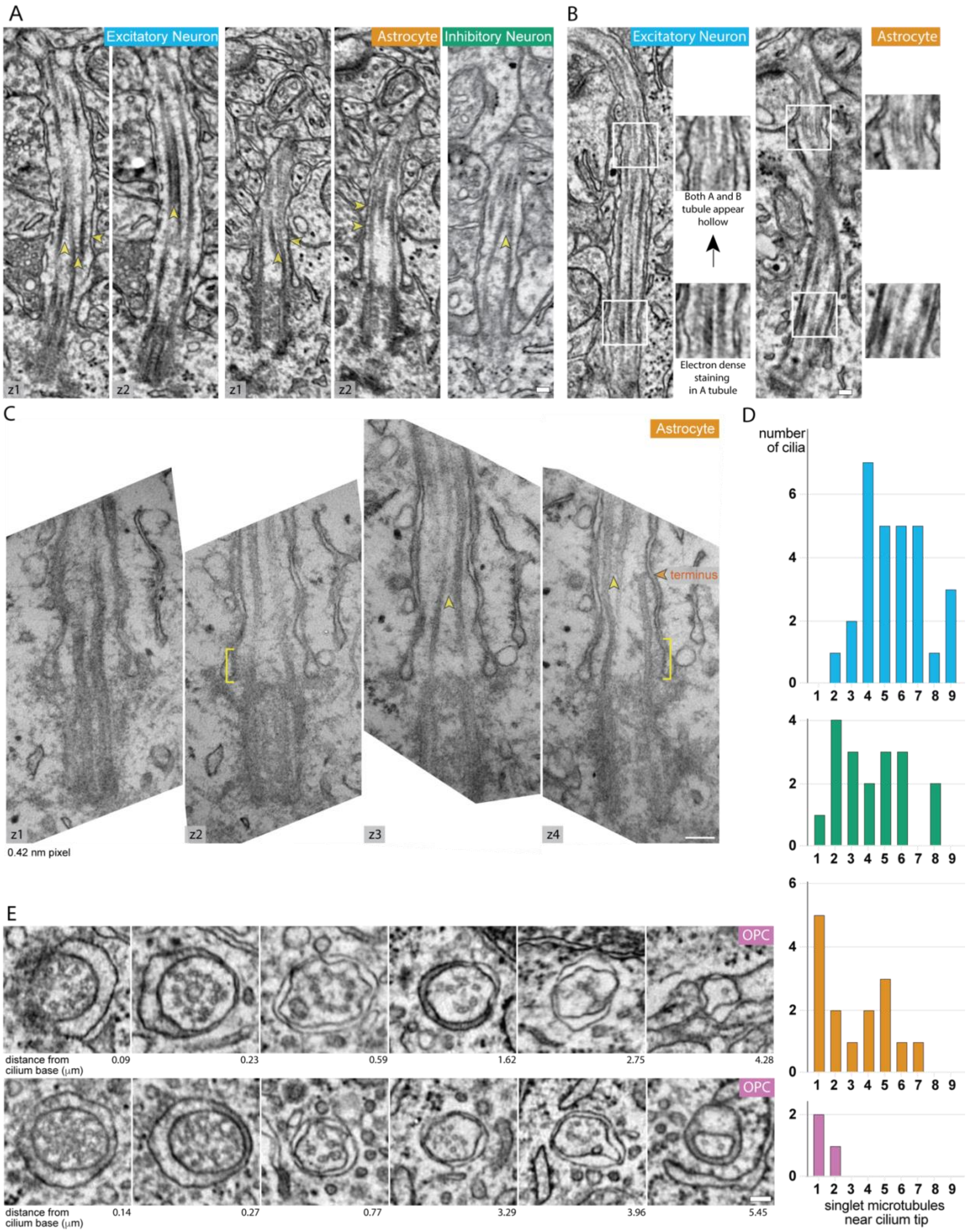


Supplemental Figure 6

**A.** Two adjacent planes each, from an excitatory neuron cilium and an astrocyte cilium, as well as a single plane of an inhibitory neuron cilium, all cut parallel to the imaging plane, show the transition of the deviant microtubule to the center of the axoneme. Densities that look like crossbridges (vertical yellow arrowheads) link the deviant filament to adjacent microtubules. The crosslinks appear to shift from one partner doublet to another as the deviant microtubule enters the center of the circle of doublets. In addition, densities between the ciliary membrane and the deviant filament (horizontal yellow arrowheads) are lost as the filament enters the center. **B.** Changes in the microtubules near the base of an excitatory neuron cilium and an astrocyte cilium cut parallel to the imaging plane. The lower inset panels show electron dense staining microtubules while the upper panels show microtubules where both walls can be resolved. **C.** The panels show the entire view of the astrocyte cilium featured in figure 3B. The yellow bracket again highlights the transition zone. In addition, densities bridging microtubules are indicated with yellow arrowheads and in the right panel, a terminating microtubule can be seen in high resolution. **D.** We evaluated the terminus of every annotated cilium in the P>270 volume and where possible, determined the number of microtubule singlets present near the cilium tip. For each cell class, the number of cilia with 1-9 microtubule singlets is graphed. **E.** Cross-sectional views along the length of two OPC cilia that were cut in cross-section show additional structures in the transition zone and an array of microtubule configurations including doublets with translucent A-tubules, singlet microtubules, and protein densities that bridge microtubules to each other and to the ciliary membrane. Scale bars = 100 nm.

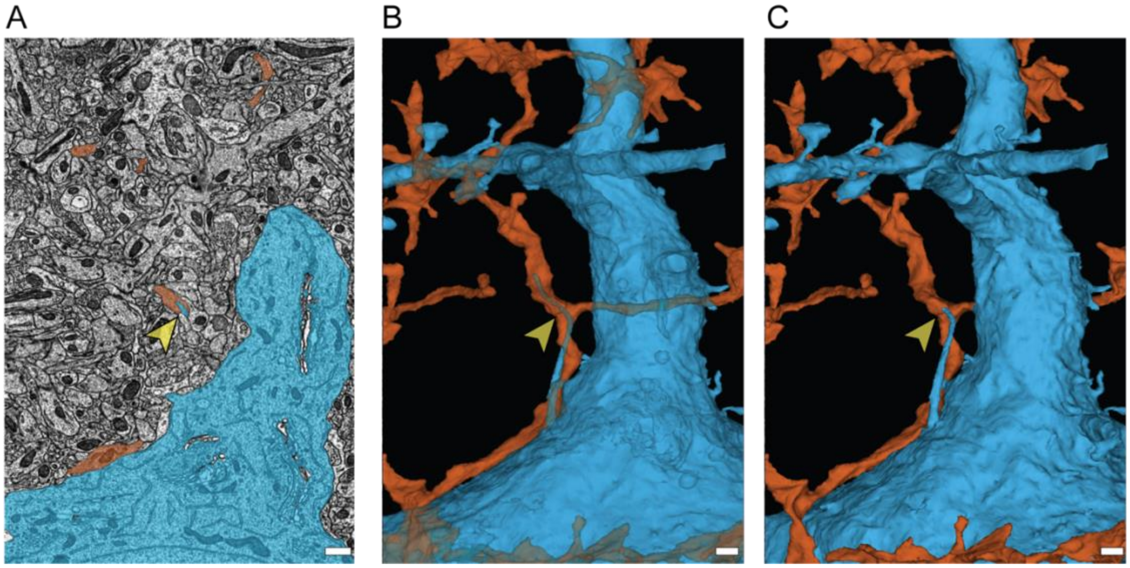


**Supplemental Figure 7** A microglial process envelops an excitatory neuron cilium

**A-C.** The single example observed of a microglial process (orange) surrounding the tip an excitatory neuron cilium (yellow arrowhead). The EM section where the view of the cilium engulfment is shown in A. A transparent and opaque 3D view of the cilium and the microglial process are shown in B and C, respectively. Scale bars are 750 nm.

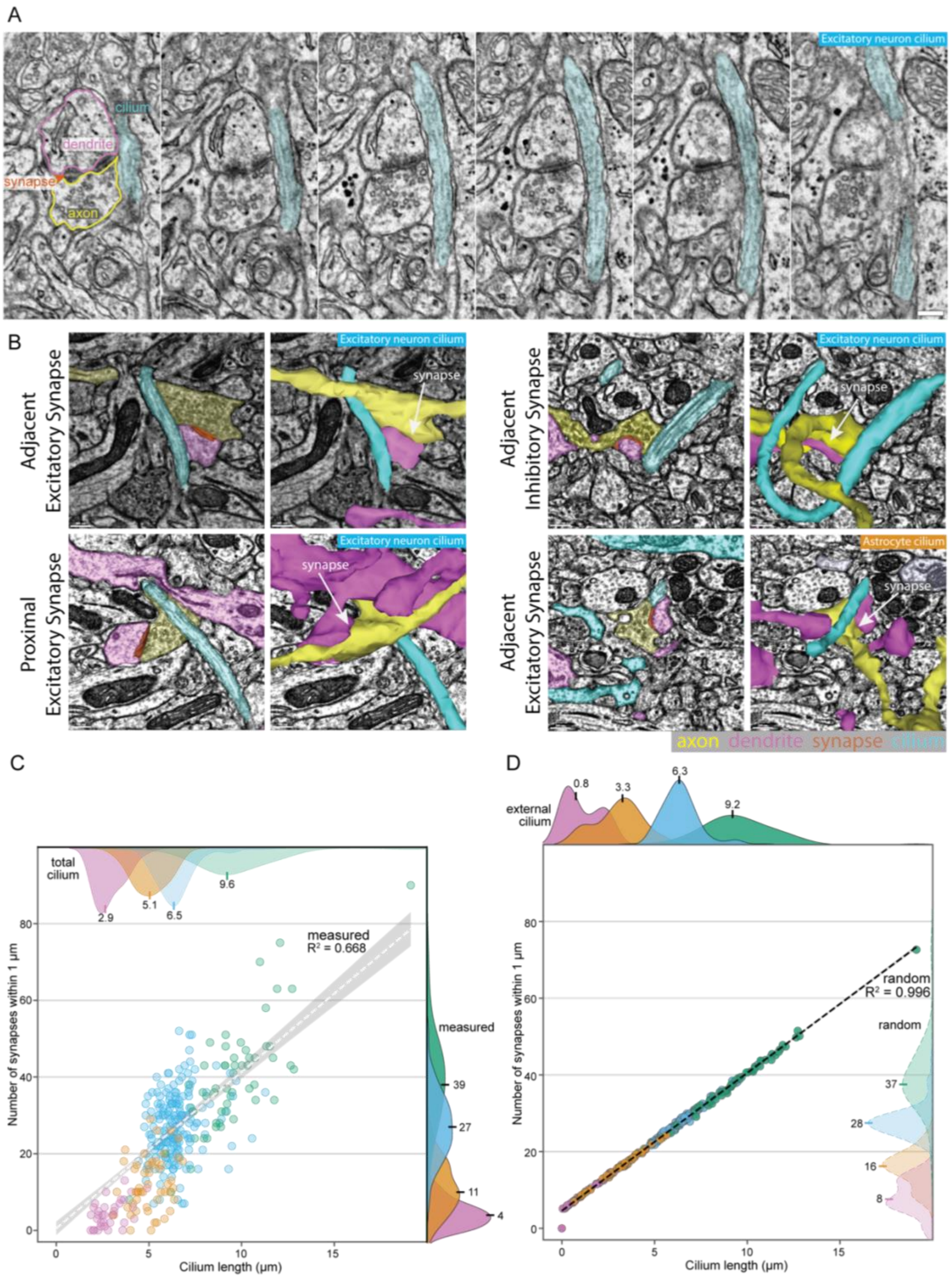


**Supplemental Figure 8: Synapses near and adjacent to cilia**

A. Serial sections of an excitatory neuron cilium that is adjacent to an excitatory synapse from the P>270 dataset. Scale bar = 200 nm. B. Four next to or near cilia are shown. For each, the left panel is a single slice of EM view of the cilium and each synapse, and the right panel includes the 3D view of the structures superimposed on the EM image. Scale bar = 300 nm. C. The number of synapses within 1 μm of each P54 cilium is graphed relative to total cilium pathlength. The line shows a linear regression fit across the dataset (β=4.91 $R^2$=0.606; shading is the 95% ci). D. The mean number of synapses within 1 μm of a meshwork of the external portion of each cilium randomly placed in 1000 positions and orientations in the EM volume is graphed as a linear regression fit (β=3.594.91 $R^2$=0.996; shading is the 95% ci). The histograms at the top show the class distributions of cilia lengths. The histograms to the right show the class distributions of the number of synapses within 1 μm.

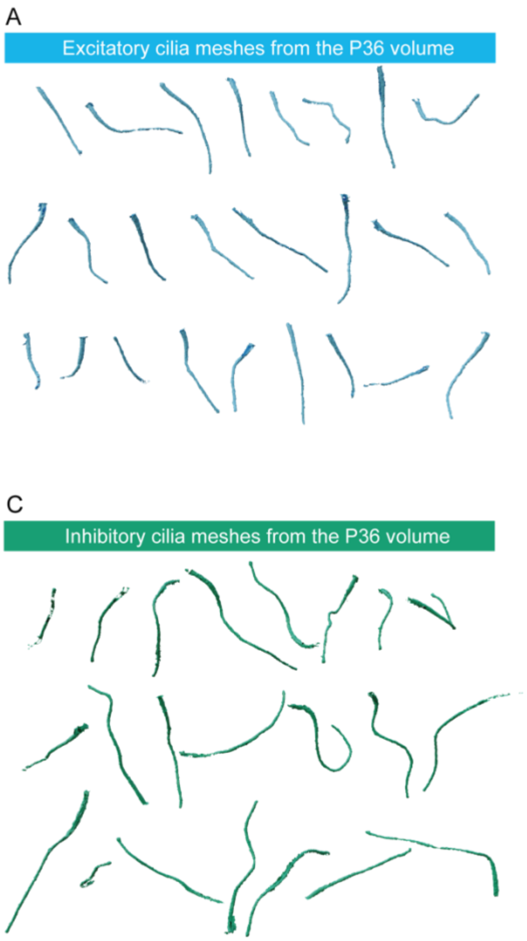


**Supplemental Figure 9: Gallery of neuronal cilia meshes**

A-D. 3D meshes of multiple cilia from the P36 and P54 volumes are arrayed. Excitatory cilia are blue and inhibitory cilia are green. The orientation of cilia in the image is representative of the orientation in the EM volume.

| Age (postnatal days) | Sex | Volume (nm) | Volume mm³ | Cortex layers | Number of cells annotated | Source |
|---|---|---|---|---|---|---|
| P36 | Male | 250 x 140 x 90 | 0.0032 | 2/3 & 4 | 385 | (Dorkenwald et al., 2022; Schneider-Mizell et al., 2021) |
| P54 | Female | 920 x 560 x 32 | 0.0165 | 2/3, 4, 5 & 6 | 521 | Allen Institute |
| P>270 | Male | 450 x 350 x 50 | 0.0079 | 1, 2/3, & 4 | 220 | (Bock et al., 2011) |

Supplemental Table 1: Features of volumetric EM datasets

**Supplemental Videos**

**Supplemental Video 1:** 3D rendering of an excitatory neuron cell and cilium

A 3D rendering of an excitatory neuron from the P36 dataset. The animation focusses in on the cilium which grows out from the soma.

**Supplemental Video 2**: Z series through both an excitatory cilium and an astrocyte cilium

The base regions of both the excitatory neuron cilium (left) and the astrocyte cilium (right) were sectioned perpendicular to the imaging plane in the P>270 dataset. Changes in the cilium diameter and the microtubule doublets as well as differences between the neuron and astrocyte cilia are illustrated by the z series. The average section thickness in the dataset is 40nm.

**Supplemental Video 3**: Z series through an excitatory neuron cilium with adjacent processes colored

This z series shows the EM of an excitatory neuron and its cilium (both are blue) with each adjacent process color coded. Dendrites are pink, axons are green and astrocytic processes are orange.

**Supplemental Video 4**: Z series through an astrocyte cilium with adjacent processes colored

This z series shows the EM of an astrocyte (orange) and its cilium (blue) with each adjacent process color coded. Dendrites are pink, axons are green and astrocytic processes are orange.

**Supplemental Video 5**: Z series a synapse adjacent to a primary cilium

The raw data z stack of a synapse adjacent to a cilium to accompany Supplemental Figure 8A
